# NOCICEPTRA: Gene and microRNA signatures and their trajectories characterizing human iPSC-derived nociceptor maturation

**DOI:** 10.1101/2021.06.07.447056

**Authors:** Maximilian Zeidler, Kai K. Kummer, Clemens L. Schöpf, Theodora Kalpachidou, Georg Kern, M. Zameel Cader, Michaela Kress

## Abstract

Nociceptors are primary afferent neurons serving the reception of acute pain but also the transit into maladaptive pain disorders. Since native human nociceptors are hardly available for mechanistic functional research, and rodent models do not necessarily mirror human pathologies in all aspects, human iPSC-derived nociceptors (iDN) offer superior advantages as a human model system. Unbiased mRNA::microRNA co-sequencing, immunofluorescence staining and qPCR validations, revealed expression trajectories as well as miRNA target spaces throughout the transition of pluripotent cells into iDNs. mRNA and miRNA candidates emerged as regulatory hubs for neurite outgrowth, synapse development and ion channel expression. The exploratory data analysis tool NOCICEPTRA is provided as a containerized platform to retrieve experimentally determined expression trajectories, and to query custom gene sets for pathway and disease enrichments. Querying NOCICEPTRA for marker genes of cortical neurogenesis revealed distinct similarities and differences for cortical and peripheral neurons. The platform provides a public domain neuroresource to exploit the entire data sets and explore miRNA and mRNA as hubs regulating human nociceptor differentiation and function.

## Background

Nociceptors are specialized primary afferent neurons that transduce and transmit painful stimuli to alert the brain of bodily harm. Their cell bodies are located in the dorsal root (DRG) or trigeminal ganglia (TG) and their developmental fate is specified by gene expression networks governed by key genes including neurotrophic receptor tyrosine kinase A (TrkA), TrkB and TrkC together with Neurogenin 1 (Ngn1) and Ngn2 (1–3).

Although comprehensive transcriptomic and proteomic analyses of mature DRGs have been performed in animal disease models and patient post-mortem tissues, human nociceptor expression signatures related to human pain disorders are scarce (4–8). Notable differences between mice and men challenge the applicability of rodent model systems for the investigation of human pain pathogenesis (9, 10). Human-cell-based model systems, such as induced pluripotent stem cells (iPSCs), which allows directed cell-type specific programming into virtually any cell type by applying specific differentiation protocols (11–15), offers the opportunity to overcome this apparent translational paresis. Human iPSCs differentiated into nociceptor like cells express fundamental nociceptor-like characteristics and emerge as a valuable tool to explore mechanisms underlying hereditary pain disorders, such as erythromelalgia or migraine, and even to develop and select personalized treatments (16–22). However, the molecular equipment of iPSC derived nociceptors and its similarity to native human nociceptors are not yet sufficiently documented, which may limit the utilization of these for mechanistic studies.

Developmental gene expression signatures and their regulation are currently only available for iPSC-derived cortical neurons (23, 24). Protein-coding genes and long non-coding RNAs (lncRNAs) have been in the focus of recent gene expression studies during cell differentiation (25, 26). At the same time, a critical role for a class of transcriptional regulators called microRNAs as hub regulators of pluripotency, balancing of cell fate decisions, as well as guiding of cell-cycle progression and self-renewal properties of human embryonic stem cells is emerging (27–34). Specific miRNA expression patterns, time trajectories and interactions with proposed target genes during nociceptor differentiation are so far unavailable but, as a decisive layer of regulatory complexity, may be highly important for neuronal cell fate decisions, morphology, target finding and finally function and dysfunction.

In order to tackle these issues, we performed unbiased mRNA::miRNA co-sequencing followed by analyses of time-course specific gene expression signatures of human iPSC-derived nociceptors (iDNs) at six different timepoints. We identified five specific sub-stages of nociceptor differentiation as well as their control by hub genes and miRNAs, highlighting the critical importance of mRNA and miRNA expression trajectories. Finally, we provide the integrative online platform NOCICEPTRA to both retrieve the presented experimental data as well as to query custom gene expression data based on specific disease and pathway enrichments during nociceptor differentiation.

## Results

### Human nociceptor differentiation involves highly correlated gene expression trajectories

Temporal gene expression patterns during the differentiation of iDNs from iPSCs were assessed at six different timepoints in three different iPSC cell lines (**Figure 1A**). Differential gene expression analysis deployed on all samples with timepoints as independent variable revealed 22’380 differentially expressed genes (DEGs). The vast majority of DEGs were protein coding genes, lncRNAs and processed pseudogenes, that showed pronounced intra-timepoint clustering as indicated by principal component analysis (PCA) and hierarchical clustering (**Figure 1B-C**; for analysis of individual cell lines see **Supplementary Figure 1A**). Enrichment analysis highlighted significantly enriched neuron-related pathways and entities indicating the transition from pluripotent cells to iDN morphology and function such as *growth cone*, *axon guidance*, *dendritic spine development*, *synapse assembly* and *glutamatergic synapse* (**Supplementary Table S1**).

**Figure 1.**
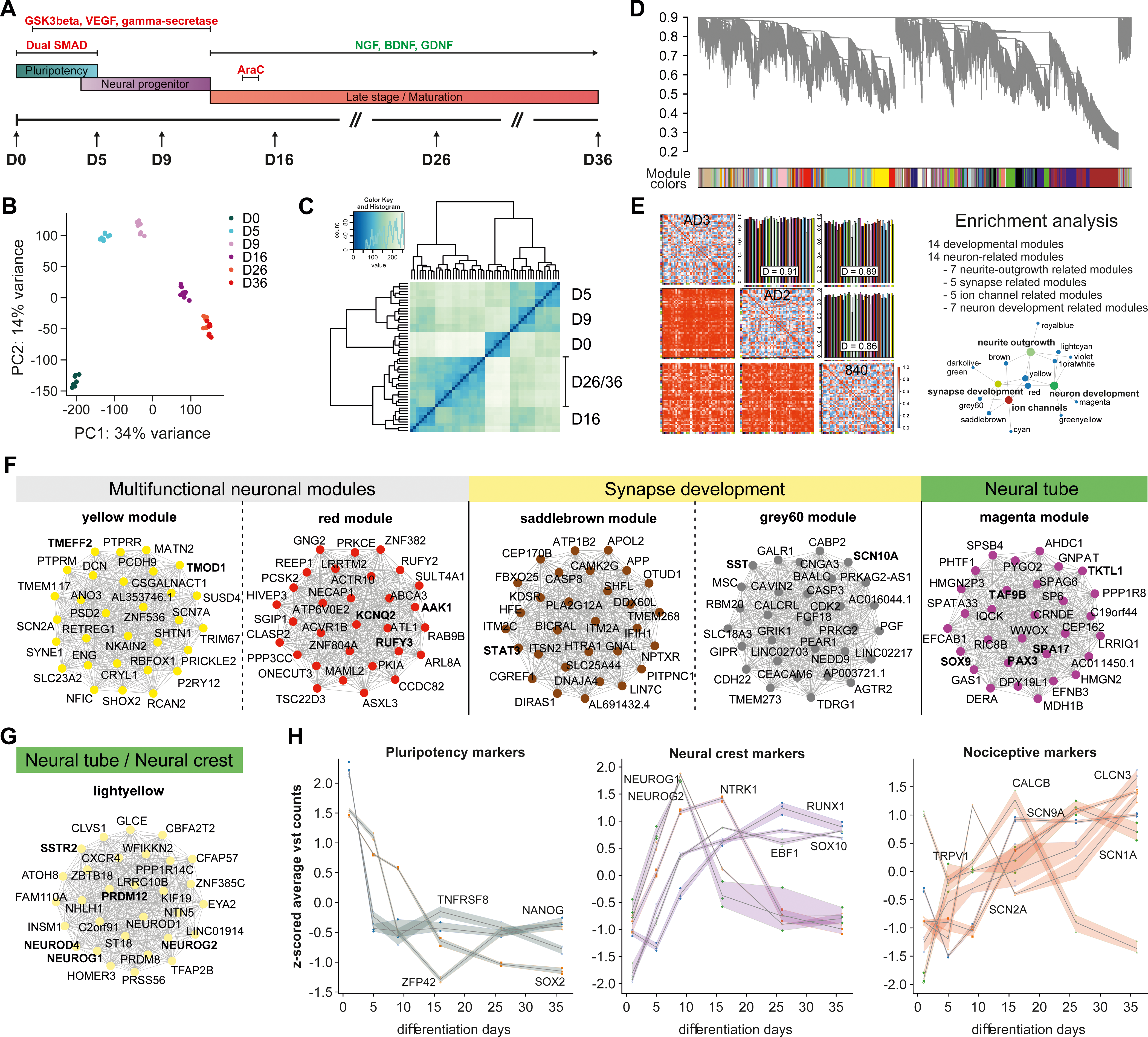
Differential gene expression analysis, time trajectories and pathway enrichments. **(A)** Protocol characteristics and timepoints for RNA harvesting based on Chambers et al (2014). **(B)** Principal component analysis and explained variance analysis of the log-normalized counts for each sample revealed high inter-timepoint specific separation and low intra-timepoint specific variation. **(C)** Overall RNA expression similarity was determined using hierarchical clustering using correlation as similarity metric. **(D)** WGCNA coexpression analysis dendrogram with module affiliations. Soft-threshold power (16) was determined assuming scale free topology. Topological overlap matrix was constructed and preservation between the three cell lines calculated and visualized in preservation as well as bar plots. **(E)** Preservation plots between the three cell lines and enrichment analysis revealed biologically conserved modules. Enrichment analysis was performed for each module applying a hierarchical best per-parent algorithm on the g:Profiler (FDR-corrected p-value < 0.05) results to obtain most specific pathways enriched, which revealed neuronal enriched modules of four entities (synapse development, ion-channel development, neurite outgrowth and development). **(F)** Top 30 hub gene networks for multifunctional neuronal modules, ion channel and synapse development related modules and neural tube development related modules. Hub-gene networks were determined using the module affiliation indicated by the k-module Eigenvector and the corresponding p-value. Hub gene networks between top 30 hub genes were constructed calculating the pairwise spearman correlations and selectively choosing a hard threshold of r > 0.7 for interconnectivity. **(G)** Top 30 hub gene network for the lightyellow module implicated in neural crest cell development **(H)** Literature-based pluripotency marker, neural crest marker and nociceptor marker time trajectories were drawn from z-scored vst-normalized counts that were averaged across the three cell lines.

Using weighted gene correlation network analysis (WGCNA) on a consensus inner joint dataset of expressed genes from all three cell lines (20’887 genes), we found 49 consensus gene modules with an average preservation of 0.89 and highly shared preservation between cell lines as indicated by the temporal trajectories (**Figure 1D-E, Supplementary Figure S1 B,C, Supplementary Figure S5-7**). Fourteen modules showed significant enrichment for neuron-related pathways associated with neurite outgrowth, synapse development, and ion channels, of which two modules (yellow and red) were highly enriched for neuronal related pathways such as synapse, neurite outgrowth and development and were therefore involved in all three aspects (**Figure 1E-F**; **Supplementary Table S2**). Networks of the top 30 hub genes of the respective consensus modules are presented based on Spearman correlations of gene expression data (**Figure 1F; Supplementary Table S3**).

Sensory neuron fate determination and neural crest cell development were initiated by two networks of highly connected genes (lightyellow module, magenta module), containing *NEUROG1*, *NEUROD2*, *NEUROD4*, *PRDM12* and *SSTR2* as significant hub-genes (**Figure 1E-F**). Analysis of important pluripotency, neural crest and nociceptor markers revealed a general time-dependent decrease of pluripotency and increase of neuronal and nociceptor markers. The neurogenic differentiation markers *NEUROG1* and *NEUROG2* followed highly correlated expression trajectories (Spearman R = 0.84, p-value < 0.05) during differentiation with peak expression at day 9, although only *NEUROG1* was abundantly expressed (∼200 TPM vs. ∼4 TPM at D9, respectively, **Figure 1H**). This finding supports successful reprogramming into small-diameter sensory neurons neurons (35, 36). *NTRK1* and *RUNX1*, two master regulators of nociceptor fate determination, showed highly correlated transient expression up to day 16, followed by an inverse-correlative expression pattern. Together with the abundant expression of the heat sensitive transducer ion channel TRPV1 and the voltage-gated sodium channel gene SCN9A, this suggested further specialization into nociceptors during maturation (**Figure 1F,H)**. In addition to RNA sequencing, we validated protein expression of SCN9A, panTRK (NTRK1, NTRK2, NTRK3), peripherin (PER), Tuj1 (TUBB3) and PIEZO2 at three different timepoints (D16-D36) during iDN differentiation, and protein expression fluorescence intensity trajectories resembled the RNA sequencing trajectories (**Supplementary Figure S2 A-B**). In accordance with the temporal RNA trajectories, the vast majority (∼ 90%) of differentiated cells were positive for panTRK at D16 and positive for SCN9A (>80%) at D36, while negative for NANOG, Ki-67, TMEM119 and GFAP (**Supplementary Figure S2C and Supplementary Figure S3 A-C)**.

The magenta consensus module likely represents the network involved in neural tube and neural crest cell development and several genes known for their contribution in neural crest/neural tube development, such as *PAX3, SOX9, TAF9B,* were identified as hub genes within this module (**Figure 1F**). Interestingly, *EFNB3* coding for the trans-synaptic organizing protein ephrin B3 was identified as a magenta module hub gene, while the roles of other hub genes within this module, such as *SPA17* or *TKTL1 (transketolase like 1)*, are largely unattended in neurons to date (**Supplementary Table S3**). *RUN* and FYVE domain containing 3 (*RUFY3*) and AP-2 associated kinase (*AAK1*) as hub genes for the red module are well known neurodevelopmental proteins and, most interestingly, *RAB7A*, prolyl 4-hydroxylase (*P4HB*) and *CDC42* as identified hub genes for protein-protein interactions are implicated in synapse development, neuronal growth and survival (37–42). Tropomodulin-1 (*TMOD1*), a tropomyosin-binding protein involved in the structuring of actin filaments in sensory neurons, and the neuronal survival factor and neuroprotectant Tomoregulin-2 (*TMEFF2*) were identified as hub genes for the yellow module (42–44). Two more modules (saddlebrown and grey60) were implicated in both synapse and ion channel development. Importantly, several sodium and potassium channels (e.g., *SCN2A*, *SCN7A, SCN10A, SCN11A and KCNQ2*) emerged as significant hub genes and may in addition to their functional roles qualify as critical indicators of neuron as well as synapse development (**Figure 1F,H**, **Supplementary Table S3-4** for top hub genes).

Since several ion channel mRNAs were expressed, functionality of iPSC-derived sensory neurons was assessed using single cell whole-cell patch-clamp recordings (**Supplementary Figure S4)**. All cells exhibited large peak inward currents indicating activation of voltage-gated sodium channels and sustained outward currents through voltage-gated potassium channels (**Supplementary Figure S4A**). Amplitudes of peak inward currents were significantly larger in AD2 cells compared to AD3 cells and this well reflects higher SCN1A mRNA expression (**Supplementary Figure S4A,E**). The resting membrane potential of ∼-55mV for AD3 cells and ∼-47mV for AD2 cellls at D36 and their capability of firing action potentials indicated a mature state of iDNs but also cell line variabilities (**Supplementary Figure S4B-D**).

Common peptidergic neuron marker expression trajectories such as TAC1 (Substance P, SP) and CALCB (CGRP2) were assessed with immunefluorescence microscopy and this revealed high expression of CALCB at D16 and TAC1 at D36 (**Supplementary Figure 1D-E, Supplementary Figure S2**).

To further support our model, we queried important markers of nociceptor development, recently determined in mice as well as transcription factors found enriched in hDRG (26, 45). Most of the developmental related markers were expressed from days 5-9 (*HES6*, *TCF4*, *EZH*, *TFAP2* and *FOXP4*), indicating their importance in human iDN differentiation (**Supplementary Figure 1G** developmental marker). Other murine nociceptor markers, such as *SYT13*, *GAL*, *TH* and *PRDM2*, peaked around days 9-16 (**Supplementary Figure 1F** nociceptor marker). All human DRG transcription factors such as DRGX, PIRT, POU4F1, RBFOX3, SKOR2 and TLX23 were expressed in iDNs and steadily increased throughout iDN differentiation with a mean TPM above 5, supporting the sensory neuron phenotype and their role in human sensory neuron differentiation (**Supplementary Figure 1H**). We further compared our iDN signatures to human native DRG and cultured human DRG data sets and found that iDNs in general were more similar to cultured human DRGs compared to native DRGs. iDNs became transcriptomically more similar to human DRGs during differentiation from D16 on, however, since DRG contain a mix of different cell types including sensory neurons, Schwann cells, macrophages and cells of the vasculature, a full overlap of gene sets cannot be expected (46, **47**), Supplementary Figures S1 J,K).

Given that many of the key genes are well known checkpoints in murine sensory neuron differentiation and human sensory neurons (26), our analysis indicates a largely conserved sensory neuron developmental program from mouse to human.

### Transcriptomic trajectories identify five distinct stages of human nociceptor differentiation

To further investigate iDN developmental stages, modules with similar gene expression time trajectories were merged by agglomerative hierarchical clustering, and singular value decomposition (SVD) was performed. Individual genes were projected onto the first two SVD-components, which resulted in a transcriptomic clock indicating different developmental stages (**Figure 2A**). Cluster analysis revealed six gene expression super-clusters with unique time trajectories that resembled *pluripotency* (2’255 genes, mainly associated with D0), *early differentiation* (6’057 genes, D0-D9), *early neural progenitor* (2’016 genes, D5-D9), *late neural progenitor* (564 genes, D16-D26), and finally the *nociceptor* stage (8’027 genes, transient expression until D36), as well as one super-cluster showing peak expression both during pluripotency and maturation (1’085 genes, D0 and D16-D36) (**Figure 2B-D**). Enrichment analysis indicative of the functional relevance for each respective stage showed that neuron-related pathways were induced as early as in the early neural progenitor phase (e.g., *neuron differentiation*). Later stages included neurite outgrowth related pathways (e.g., *neuron projection*) and synapse-related pathways (e.g., *glutamatergic synapse*, *synaptic membrane;* **Figure 2E, Supplementary Table S5**) and several neuronal related diseases, such as *cognitive disorders*, *neurodegenerative diseas*e and *nervous system disease* (**Supplementary Table S6**). We further identified hub genes for each differentiation stage based on functional connectivity using StringDB network analysis. This revealed *MYC* and also *RHOA* as hub regulators in the pluripotency phase, and *AKT1* and *SRC* encoding serine/threonine kinases as well as *UBC* in the nociceptor stage (**Supplementary Table S7**). We found that *SOX9* (magenta module) and *MED13* (grey60 module), which are strongly involved in neural tube/neural crest cell development, were most abundantly expressed between days 5-9, (**Figure 1F**, **Supplementary Table S3,7**).

**Figure 2.**
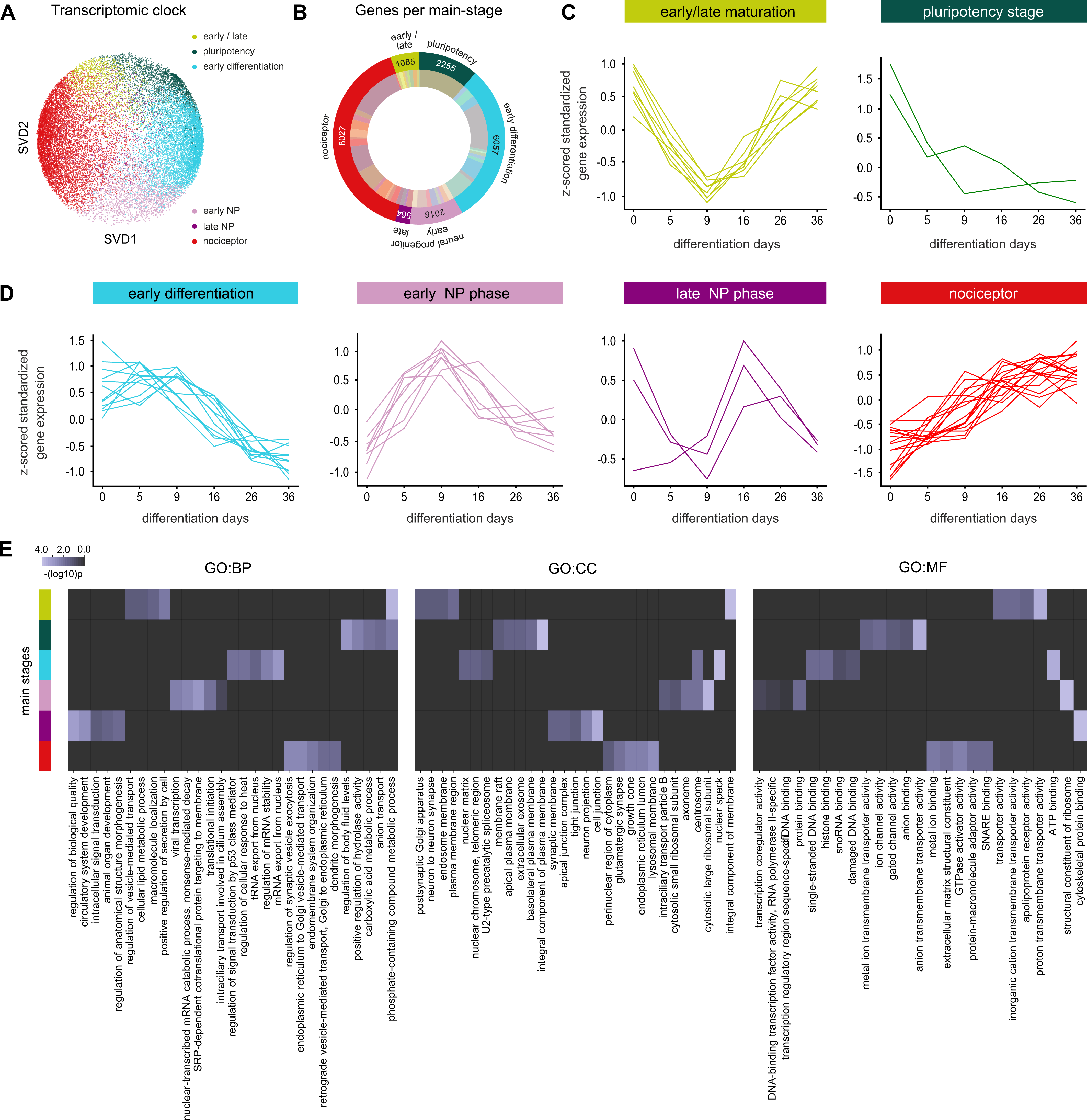
mRNA trajectory-based stages of nociceptor differentiation. **(A)** Singular value decomposition (composition of three matrices) performed along the gene expression axis using averaged z-scored vst-normalized counts and projected onto the first two components revealed a transcriptomic clock indicative of different nociceptor stages. Dot color represents the clusters affiliation identified by merging WGCNA module with similar expression profiles via hierarchical clustering using the “average” agglomerative algorithm and mean z-scored variance-stabilized counts. **(B)** Pie-chart of the distribution of mRNAs for each stage of nociceptor differentiation with number of genes. Outer circle represents the main stages during nociceptor development and inner circle shows the number of WGCNA modules affiliated with each main stage as well as the overall number of genes associated with each main stage and module. **(C-D)** Mean z-scored vst-normalized curves for each module affiliated with the representative main-stage. **(E)** Top 3 g:Profiler enrichments for each main stage, ranked based on the negative log10 p-value for 3 annotation spaces (GO:BP, GO:CC, GO:MF). The best-per parent approach was used to identify specific enriched pathways as described in *Zeidler et al. (2020)*.

### microRNAs show differentiation stage-specific expression

As numerous protein-coding genes are regulated by miRNAs, we performed small RNA co-sequencing from the same samples to identify timepoint specific expression trajectories of differentially expressed miRNAs (DEmiRs) during nociceptor differentiation. PCA and hierarchical clustering deployed on all samples revealed 979 DEmiRs that similarly to DEGs showed pronounced intra-timepoint clustering although with slightly higher variances (**Figure 3A-B**; for analysis of individual cell lines see **Supplementary Figure 8A-C**). WGCNA analysis of a consensus inner joint miRNA dataset from all three cell lines (1’647 miRNAs) revealed 17 highly correlated miRNA consensus modules as indicated by temporal trajectories with an average preservation of 0.88 (**Supplementary Figure S8E, Supplementary Figure S9**). SVD resulted in five main differentiation stages based on miRNA expression trajectories that overall resembled the gene expression trajectories (**Figure 3C-E**). Interestingly, no miRNA trajectory cluster was found that resembled the pluripotency/maturation stage. The top regulated miRNAs per super-cluster were hsa-miR-302a-5p for the pluripotency stage, hsa-miR-25-3p for the early differentiation stage, hsa-miR-363-3p and hsa-miR-96-5p for early and late neural progenitor stage, and hsa-mir-183-5p for the nociceptor stage (**Figure 3F**). miRNA qPCR validation of hsa-miR-302a-5p, hsa-miR-25-3p, hsa-miR-103a-3p, hsa-miR-355-5p and hsa-miR-146-5p revealed similar time-dependent trajectories as found mRNA, miRNAs were choosen based on temporal trajectories as well as expression strength to cover low expressed (hsa-miR-103a-3p) but also highly expressed miRNAs (hsa-miR-302a-5p, **Supplementary Figure 8F**).

**Figure 3.**
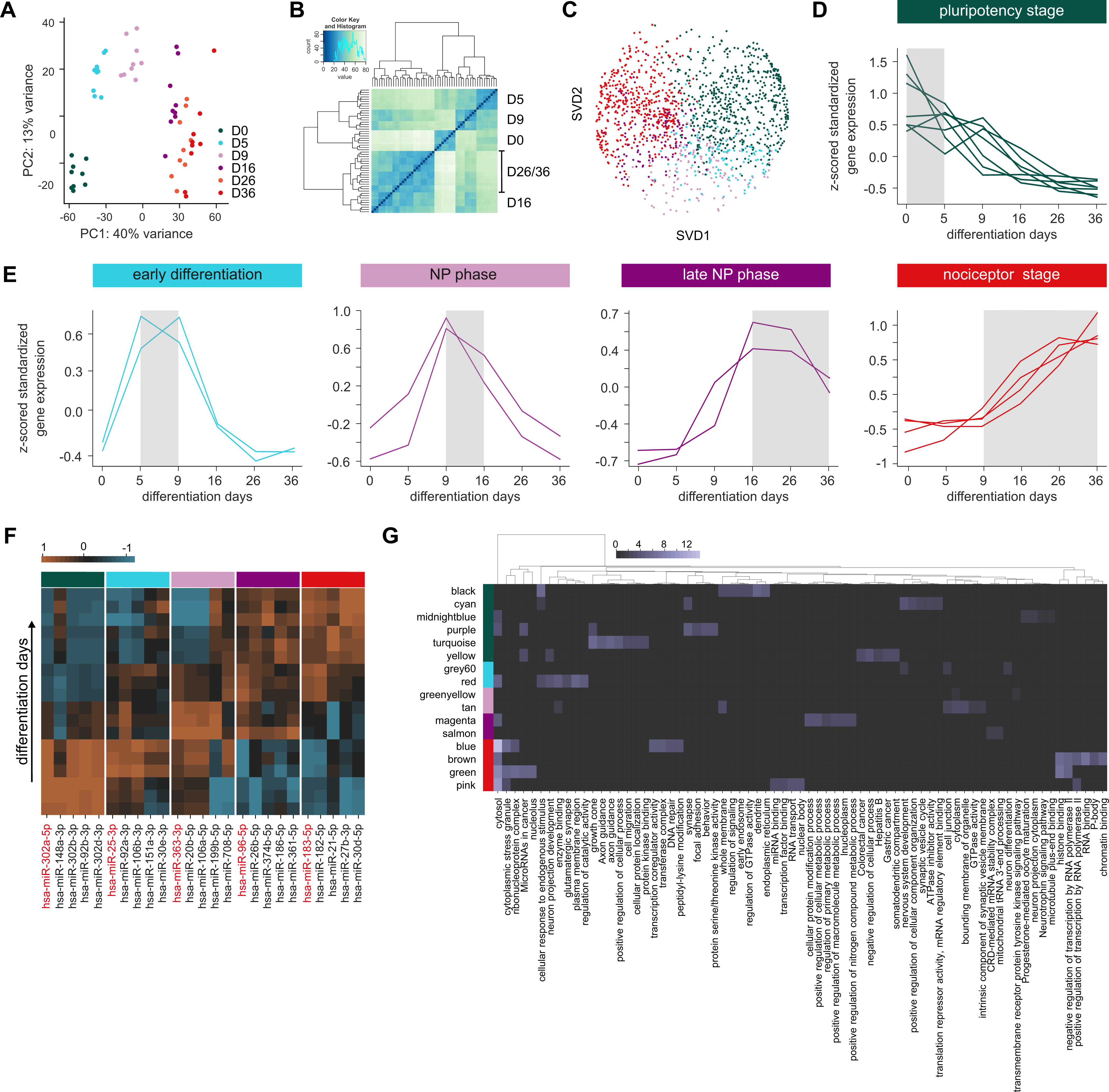
Differential miRNA expression and trajectory-based differentiation stages. **(A)** Principal component analysis along the sample axis of miRNA expression during iDN development projected onto the first two principal components. Explained variance for each principal component (PC1, PC2) is annotated at the corresponding axis and inter-timepoints as well as inter-cell line variance investigated. **(B)** Hierarchical clustering using the “average” agglomerative algorithm based on similar miRNA expression revealed high intra-timepoint clustering with low inter cell type variances. **(C)** Singular value decomposition (SVD) of each individual miRNA based on expression projected onto the first two SVD components. Each individual miRNA is depicted by a dot and colored by the cluster affiliation determined applying the “average” hierarchical clustering algorithm on the main averaged temporal WGCNA module trajectories, which revealed a transcriptomic clock of miRNA expression with 5 main stages modelling the differentiation of iPSC-derived nociceptors. **(D-E)** Resulting clusters were ranked based on the transcriptomic clock and average (cell line) z-score vst-normalized expression patterns representing time trajectories of each individual WGCNA module belonging to the main stage and are visualized with line plots. **(F)** miRNA abundance analysis for each individual differentiation stage based on baseMean expression (normalized DEseq2 counts) showing the Top 5 most abundant miRNAs for each main stage and the temporal expression patterns were depicted in a heatmap (z-score standardized). **(G)** Reverse miRNA enrichment profiling for each WGCNA module ranked according to the main differentiation stages was determined using the so constructed miRNA_edge.csv database target-score. Only miRNA::gene interactions with a target-score above 70 (most of them highly predicted and validated) and an inverse correlation below r < -0.7 were considered and top regulated pathways of targeted genes were determined using g:Profiler and ranked by the negative log10 p-value.

The majority of the consensus miRNA modules was affiliated with the pluripotency stage (7 modules) and the nociceptor phase (4 modules), indicating that the miRNAs within these modules may function as critically important master switches controlling large gene sets essential for these stages (**Figure 3G**).

To investigate which pathways were regulated by miRNAs during iDN development, a miRNA::mRNA target edge database was constructed that included a combined target score based on the DIANA microT-CDS prediction score, the correlation between miRNA and mRNA expression (vst counts) in our samples, the percentile-ranked miRNA base expression as well as the target validation status (using StarBase, miRTarBase and Tarbase). All targeted genes (correlation < -0.7, target-score > 70) for each module were determined and reverse enrichment analysis was performed. Predominantly miRNA modules belonging to the pluripotency and early differentiation phases were enriched for neuron related pathways such as *growth-cone*, *axon guidance* (*turquoise module*, pluripotency phase), *synapse* (*purple module*, pluripotency phase) and *glutamatergic synapse* (*red module*, early differentiation phase). In particular, the miR-17∼92 cluster was highly expressed and associated with the early differentiation of iPSC-derived nociceptors (D5-D9). In addition, hsa-miR-20a-5p, hsa-miR-92a, hsa-miR-19b and hsa-miR-18a were highly regulated and associated with glutamatergic synapse formation, neuronal cell body and endomembrane system organization pathway enrichments (**Figure 3F-G**).

### miRNAs regulating hub genes

Since hub genes (kME > 0.8) were the main regulators per module, only those were considered for miRNA::mRNA target analysis. An adjacency matrix was calculated and UMAP dimensional reduction on a sparse target-score matrix followed by k-Means cluster analysis (n clusters = 8) performed to identify miRNA::mRNA interaction clusters associated with nociceptor differentiation stage (**Figure 4A**). Cluster analysis revealed four miRNA::mRNA interaction clusters associated with the nociceptor stage (**Figure 4A**), which were significantly enriched for neuron-related pathways such as *axon guidance* (cluster 7)*, synapse* (cluster 1), *presynapse* (cluster 2), and *glutamatergic synapse* (cluster 6) (**Supplementary Table S8**).

**Figure 4.**
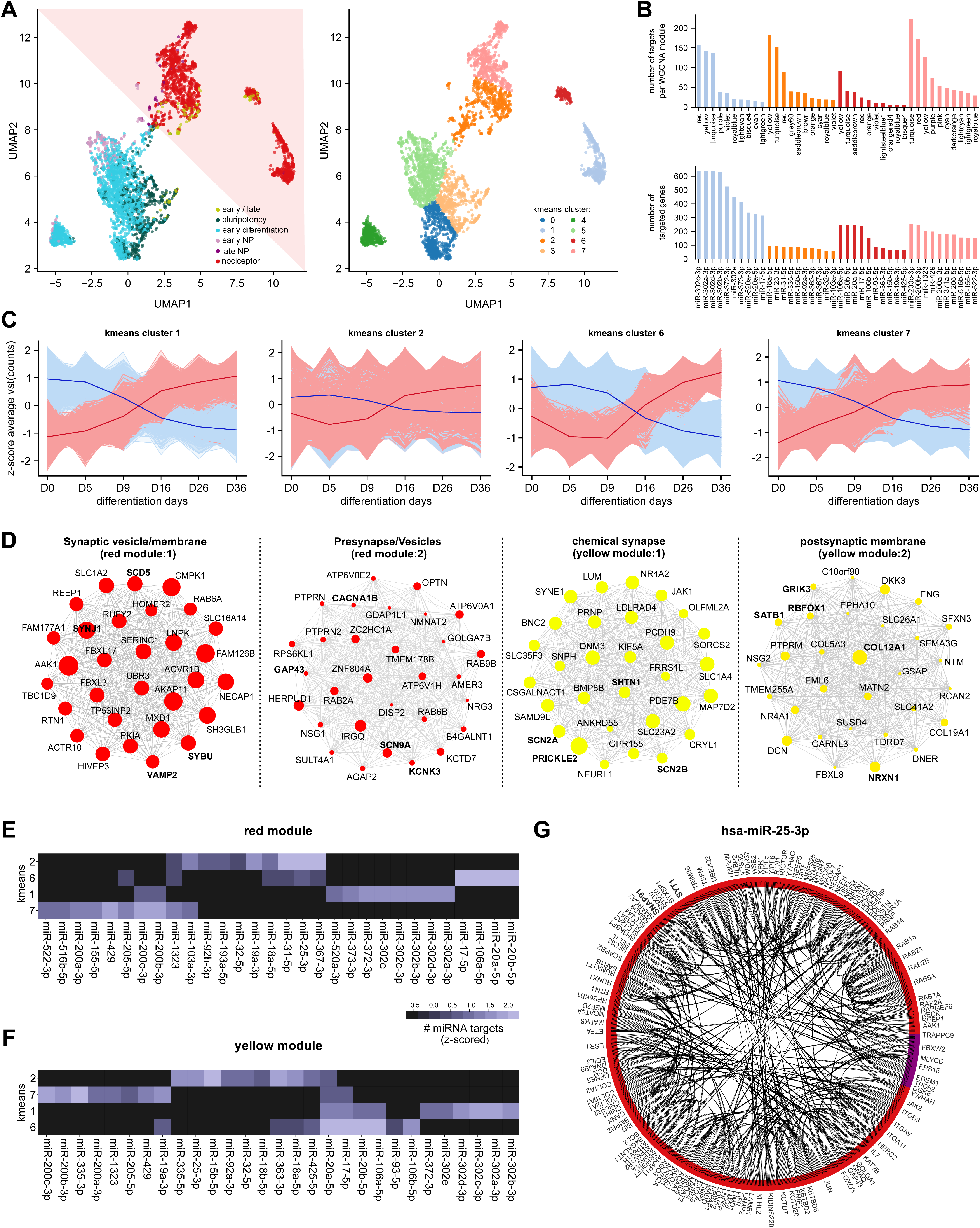
miRNAs regulate pre- and post-synapse development via module hub genes. **(A)** A miRNA::hub-gene target-space interaction map (adjacency matrix using the target-score between each miRNA::gene interaction) was constructed by using the miRNA_edge.csv database with a target-score above 70 and an inverse correlation below -0.7 between the gene target and the miRNA. UMAP dimensional reduction was performed to reduce the resulting adjacency matrix along the targeted genes axis, which revealed higher local connectivity of genes targeted by the same miRNAs as well as high local connectivity of genes representing the same main stage. Each individual dot following UMAP construction was colored according to the affiliation to the differentiation main stage, which revealed separation between early and late iDN differentiation based on miRNA targeting. k-Means clustering (with n_cluster = 8) of the adjacency matrix revealed 8 clusters of targeted genes targeted by different miRNA groups. K-Means affiliation is projected onto the UMAP and k-mean clusters affiliated with late-stage nociceptor development were chosen further analysis. **(B)** k-Means clustering revealed 4 clusters corresponding to the late stages of iDN differentiation, WGCNA module affiliation for each gene was determined and the number of targets within each WGCNA module for each k-Means cluster was determined as well as the number of genes targeted by each individual miRNA. Results were ranked descending to evaluate miRNA as well as module importance for each k-Means cluster. **(C)** 4 clusters showed neuronal enriched gene pathways in the g:Profiler analysis; miRNA (lightred) and gene expression (skyblue) trajectories for each of the k-Means clusters was represented as z-scored vst-normalized temporal expression of each gene and miRNA trajectory belonging to the k-Means cluster; an averaged gene (red) and miRNA (blue) expression curve was calculated and Pearson correlation calculated between the averaged curves. **(D)** Top 30 miRNA targeted hub gene networks for each WGCNA module per k-Means cluster was constructed using the number of miRNA interactors as node-size as indicator of suppression strength within the module. **(E-F)** miRNAs with the highest number of miRNA targets using groupby and counting functions provided by python for the red and yellow modules were determined for each k-Means cluster and z-scored data visualized for each of the 4 clusters. **(G)** Protein-protein interaction network (StringDB) of all hsa-miR-25-3p targets and the main differentiation stage affiliation indicated by the arc color visualized using a chord interaction plot.

Of the consensus gene modules, especially the red, yellow, turquoise, violet and purple modules were highly targeted by miRNAs belonging to the miR-302 family, the miR-17∼92 cluster, the miR-106∼25 cluster, hsa-miR-31 and the miR-200 family (**Figure 4B**). The anti-correlative miRNA::mRNA expression trajectories revealed two distinct patterns: clusters 1 and 7 showed a transient increase in gene expression and decrease in miRNA expression, whereas clusters 2 and 6 showed no change until D9, followed by a sustained increase in gene expression and decrease in miRNA expression (**Figure 4C**). This documents the critical importance of miRNA inhibitory control over certain gene sets, which is released by decreasing miRNA expession during iDN development.

Overall, 586 miRNAs were highly expressed during pluripotency and expression decreased with proceeding differentiation. To further dissect the role of specific miRNAs in the development of iDNs, we performed functional enrichment analysis of the hub-genes of the consensus gene modules in the same interaction cluster, which are therefore targeted by similar miRNAs. This revealed a strong post-transcriptional control of synapse-related pathways (mainly in the red and yellow consensus gene modules, **Figure 4D, Supplementary Table S9**). Within the red module, several hub genes targeted by the miR-302 family were significantly enriched for synaptic vesicle related pathways (VAMP2, *SCD5*, *SYBU*, *SYNJ1,* cluster 1). Hub genes that were predominantly targeted by hsa-miR-367-3p, hsa-miR-25-3p and hsa-miR-31-5p were enriched for presynapse/ion-channel complexes (*SCN9A, KCNK3, GAP43, CACNA1B*, cluster 2) (**Figure 4D-E, Supplementary Table S9**). In contrast, miRNAs of the miR-302 family, hsa-miR-25-3p, and hsa-miR-363-3p targeting genes within the yellow module, were more implicated in the regulation of chemical synapse transmission (*SCN2A, SHTN1 and PRICKLE2, cluster 1*) and postsynapse (GRIK, RBFOX1, *COL12A1, NRXN1 and SATB1*, cluster 2, **Figure 4D,F**).

Several miRNAs such as hsa-miR-25-3p targeted genes within several clusters and modules and could therefore serve as master switches suppressing genes associated with mature neuron machinery and function (**Figure 4D-F**). Enrichment analysis of all high confidence targets revealed an important function in nervous system and postsynapse development (**Supplementary Table S10**). Furthermore, the synaptic genes *SYT1* and *SNAP91*, which are associated with synapse development but also implicated in central nervous system disorders such as autism spectrum disorder or epilepsy, emerged as high-confidence targets of hsa-miR-25-3p (**Figure 4G**) highlighting this miRNA as an interesting novel target for neuron-related disorders.

### Neurite outgrowth modulated by miRNAs

The formation and outgrowth of fine processes such as dendrites and axons is a prerequisite to connect sensory neurons to their target tissue and to postsynaptic neuronal assemblies (48), but also to allow for nerve regeneration following peripheral nerve injury (49). Of the seven consensus gene modules related to neurite outgrowth (**Figure 1E**), three (royalblue, violett and floralwhite) were strongly enriched for neurite outgrowth, cell migration and cytoskeleton regulation (**Figure 5A,B**, **Supplementary Table S2**). All three modules showed a strong increase in gene expression after D9 and contained well known hub genes that are mechanistically involved in neuronal outgrowth (royalblue: *COL1A1, MCAM, RHOC, RHOB, RHOJ, DUSP6*; violet: *TFBR1, FAM114A1, ROCK2, POSTN*, floralwhite: *AMIGO1, CLSTN1, PHACTR1, MAPK1, PTPN1*; **Figure 5A-B, Supplementary Table S4**).

**Figure 5.**
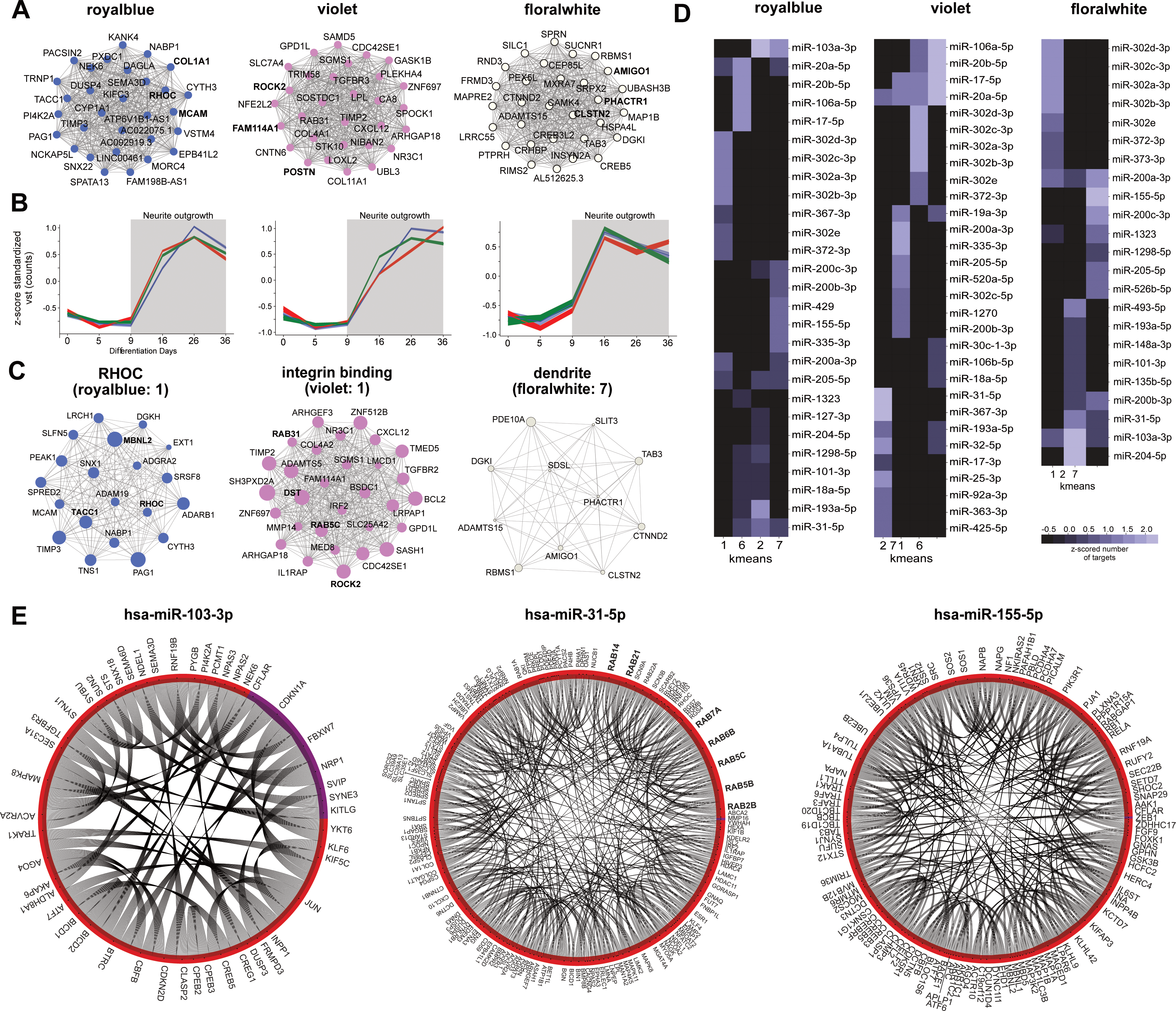
Neurite outgrowth related miRNA::gene interactions. **(A-B)** Top 30 hub gene networks (edges represents gene::gene correlations) based on the ranked kME for the royalblue, violet and floralwhite modules as well as the averaged standardized gene expression trajectories (vst(counts)) for all three cell lines and modules highly correlates with neurite outgrowth. **(C)** Number of top hub genes regulated by miRNAs predominantly targeting k-Means cluster 1 and 2 for the royalblue and violet module; clusters were named according the G:Profiler enrichments. **(D)** miRNAs with the highest number of targets for the royalblue and violet module were determined for each k-Means cluster and hierarchical clustering along the z-scored cluster axis performed. **(E)** Chord network analysis of hsa-miR-103a-3p, hsa-miR-31-5p and hsa-miR-155-3p targets with edges representing StringDB interactions and arc colors representing the main differentiation stage affiliation of the genes.

Based on their expression trajectories and target prediction scores emerging from our pipeline, we selected several interesting miRNAs that could be critically involved in these processes as regulators of larger gene sets spanning multiple clusters and modules. Hub genes such as *RHOC, MBNL2, TACC1* from the royalblue and *ROCK2, DST* and *RAB’s* from the violet module were predominantly targeted by the human miR-302 family (cluster 1, **Figure 5C-D**). The resulting regulation of RHO GTPase activity and integrin binding putatively suppressed neurite outgrowth and cell migration until D5. Since the miR-302 family shows a diverse target-spectrum, an analysis of all high-confidence targets (target-score > 0.75, r < -0.7) was performed to pinpoint functionality even further. This revealed a significant enrichment for neurite outgrowth related pathways *microtubule binding*, *axonal transport, growth-cone* and *dendrite* (**Supplementary Table S11**).

Interestingly, the hsa-miR-103a-3p target space within cluster 7 appeared to be implicated in the suppression of neurite outgrowth and axonal elongation by targeting genes belonging to the floralwhite *dendrite* module but also to clusters enriched for *phosphatidylinositol-3- phosphat binding* (violet) and *cell morphogenesis* (royalblue) (**Figure 5 C-E, Supplementary Table S12**).

For a third relevant miRNA, hsa-miR-31-5p, high-confidence targets were found in all 3 modules in most interaction clusters, suggesting a general control via this miRNA and therefore an important role in neurite outgrowth and axonal development (**Figure 5E, supplementary Table S13**). When performing pathway enrichment for all anti-correlated high-confidence targets, it became evident that hsa-miR-31-5p might further exert its function by suppressing expression of RAB GTPases *RAB14, RAB1A, RAB21, RAB2B* (**Figure 5D,E**), which regulate membrane trafficking (50). We further identified hsa-miR-155-5p as significantly enriched for neurite outgrowth/synapse related pathway predominantly targeting hub-genes of cluster 7, which is crucial for the suppression of axonal elongation in mice (51) (**Figure 5D,E, Supplementary Table S14**).

### Ion channel expression as a signature of functional iDN maturation

Ion channels are essential for the general functioning of cells and in particular of excitable mature neurons including nociceptors. Therefore, we investigated the expression trajectories of all currently assigned ion channels (**Figure 6A**). Six consensus gene modules were significantly associated with ion channel activity, including the brown and cyan modules (**Figure 6B**; *brown*: top hub genes: *ESRP1, KCNK6*; enrichments: gated channel activity, *integral component of the plasma membrane*; *cyan*: top hub genes: *AL365203.2, CYB561D1, SLCO2B1, NGFR, BIN1*; enrichments: *plasma membrane, ion channel regulator activity,* **Supplementary Table S2**). *NGFR* and *BIN1* are well established regulators of ion channel trafficking to the membrane and both were highly correlated with the expression of the DRG specific voltage-gated sodium channels *SCN10A* and *SCN11A* (r > 0.86, p < 0.05) (52).

**Figure 6.**
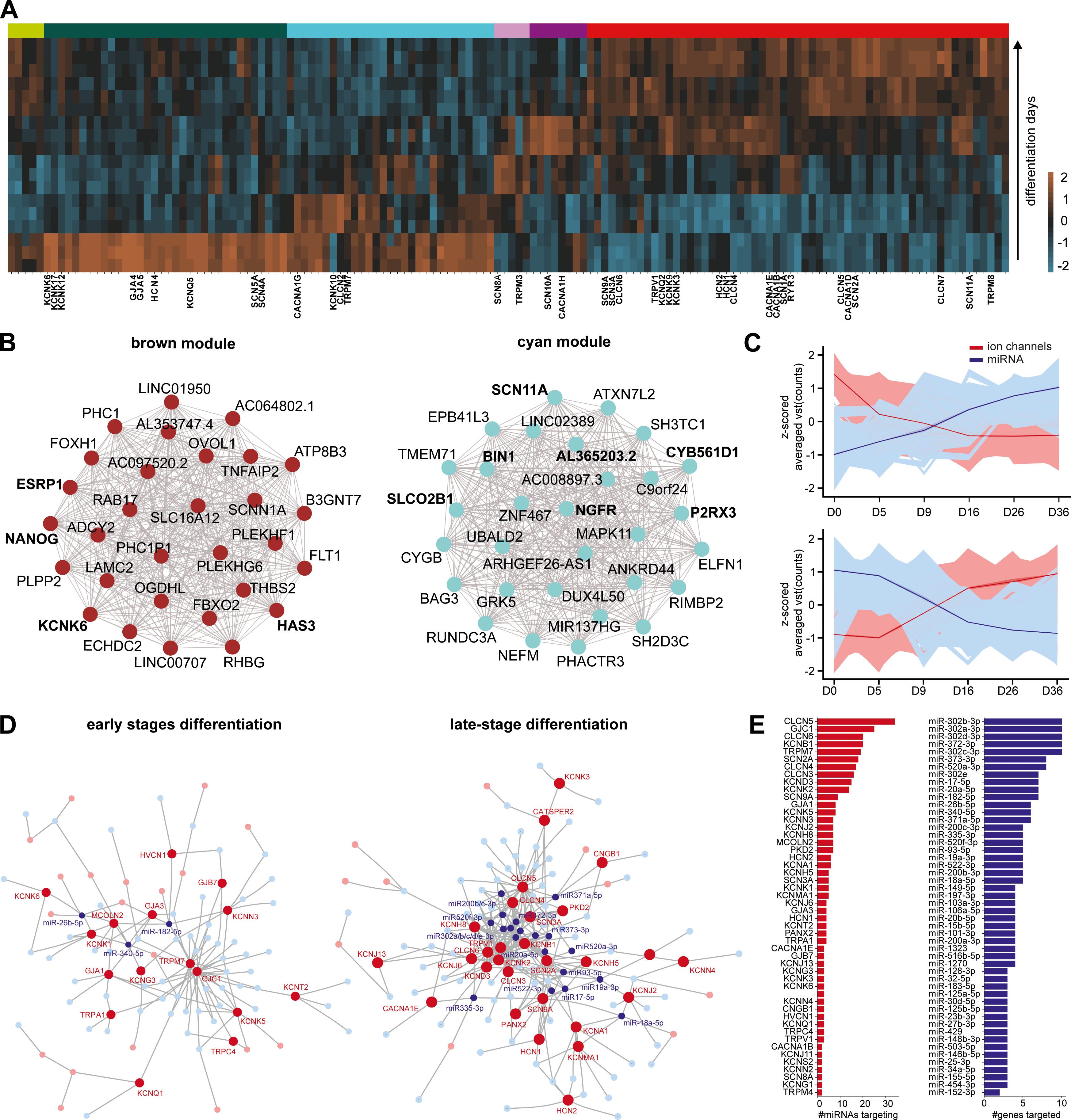
miRNAs regulate ion channels via hub genes. **(A)** Heatmap of ion channel expression during iDN development based on vst-normalized counts averaged across the cell lines, ranked based on main differentiation stage affiliation and iDN developmental progress was calculated. **(B)** Top 30 hub genes of the brown module highly enriched for ion channel related gene pathways and expressed in iPSC-stage and Top 30 hub genes of the cyan module with ion channels marked in red as top hub regulators. **(C)** Time trajectories of ion channels (blue) expressed in the early stages and late stages of iDN differentiation and the mean projection curve, as well as miRNA expression trajectories (rose) found to be regulating ion channels (right, top). **(D)** miRNA::mRNA network analysis considering the target-score as edge weight, miRNAs (blue) and genes (red) with more than 3 edges are annotated and node size represents the eigenvector centrality **(E)** The number of genes targeted by a microRNA (red barchart) and the number of targeted genes by miRNAs (blue barchart) represented in ascending orders.

A large number of ion channels was already expressed at the iPSC stage and early differentiation stages, and underwent a drastic reduction of expression during differentiation (**Figure 6A**). This included potassium channels *KCNQ1, KCNQ5, KCNS1 and KCNK6,* voltage-gated sodium channels *SCN4A, SCN5A* as well as *HCN4*, *CLCN2, CACNA1G* and *TRPM7*. As expected, several ion channels were predominantly expressed in the nociceptor stage (e.g., *SCN1A, SCN2A, SCN3A, SCN9A; KCNQ2, KCNJ6, KCNJ13, KCNJ16; CLCN3, CLCN4; HCN1, HCN2*)

(**Figure 6A)**. The two hub genes *SCN10A* (*grey60 module*) and *SCN11A* (*cyan module*) peaked around D16, and *SCN8A* a voltage-gated sodium channel which is essential for neuron function in mammalian nerve tissues and in the pathogenesis of neuropathic pain (53–55) peaked already at D9 at the transition from pluripotency to neuronal differentiation (**Figure 1F, Figure 6B**). This suggests a critical role of *SCN8A*, *SCN10A* and *SCN11A* not only for nociceptor function but also differentiation and maturation. In general, these findings support the idea that neuronal excitability and action potential discharge may not only be required for neuronal function, but also critically affect cell fate decisions and establishment of target tissue innervation and neuronal circuitry (56).

### miRNAs regulating ion channel expression

Analysis of miRNAs targeting ion channels revealed two highly connected communities of miRNA::ion channel networks that could be separated based on their differentiation timepoint affiliation. Ion channels and corresponding miRNAs revealed highly significant inversely correlated trajectories early (D0-D5, **Figure 6C**) and later during differentiation (D16-D36; **Figure 6C**), as indicated by the mean trajectories drawn for miRNA and mRNA vst counts. The gap junction related ion channels *GJC1* and *GJA1* were among the most targeted ion channels in the iPSC stage (**Figure 6D,E**). Also, the transient receptor potential-melastatin-like 7 (*TRPM7*) was expressed mostly in the iPSC stage, and appeared to be post-transcriptionally controlled by at least 17 miRNAs during differentiation (**Figure 6D,E**). At the more mature stages, several ion channels emerged, whose relevance for the transduction of painful stimuli, excitability or action potential propagation is well accepted. Two important nociceptor ion channels *SCN2A* and *SCN9A* (57, 58) were highly and inversely correlated with hsa-miR-20b-5p, hsa-miR-15b-5, hsa-miR-18-5p, hsa-miR-423-3p, hsa-miR-93-5p and others, which decreased during the differentiation, while *SCN2A* and *SCN9A* expression increased. Also, the chloride transporters and channels *CLCN4-6* were strongly targeted by miRNAs, with *CLCN6* being controlled by 14 miRNAs and *CLCN5* being targeted by 23 miRNAs. Interestingly, all three chloride channels were targeted by members of the hsa-miR-302 family. In general, the miR-302 family and hsa-miR-372 was found to suppress more than 10 ion-channels (**Figure 6E**).

### NOCICEPTRA resource

For general use and utilization of our data set, we provide the online tool NOCICEPTRA based on the Python “*streamlit*” framework, to analyze and visualize time trajectories of genes and miRNAs of interests. Both DESeq2 variance-stabilized counts as well as TPMs can be queried for each gene and miRNA present in the dataset, and a correlation matrix for the queried genes calculated. We further implemented an integrated exploratory data analysis tool, to browse all consensus gene and miRNA modules, enrichments, and the top 30 hub gene regulators, as well as miRNA interactions with the hub regulators. In addition, KEGG (www.genome.jp/kegg) and disease pathways (www.disgenet.org) can be browsed, and standardized Pearson residuals calculated to identify potential windows of susceptibility during iDN differentiation, together with an integrated g:Profiler enrichment analysis (www.biit.cs.ut.ee/gprofiler) and StringDB PPI analysis (http://www.string-db.org/, **Figure 7A**). Also, single miRNA target spaces can be queried with adjustable parameters (e.g., target-score, correlation) to determine the putative function of miRNA during iDN differentiation indicated by StringDB PPIs and enriched pathways (**Figure 7A**) and single genes or gene sets miRNA target spectra determined.

**Figure 7.**
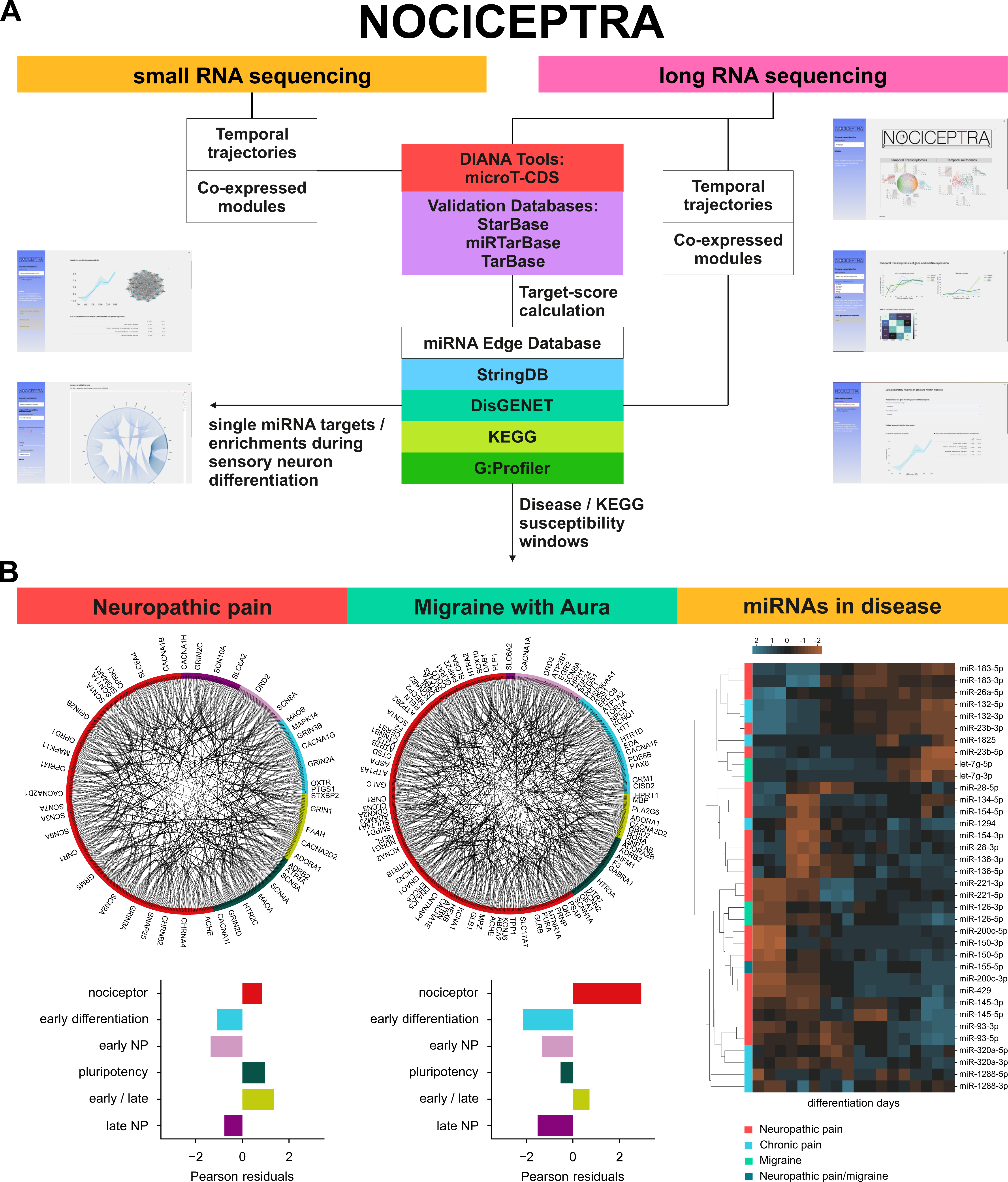
The NOCICEPTRA tool and disease enrichment analysis. **(A)** Schematic diagram of all databases incorporated in the NOCICEPTRA tool as well as analysis outputs. **(B)** Susceptibility windows of diseases determined using the NOCICEPTRA tool with OpenTarget.com risk associations of migraine with aura and neuropathic pain. Contingency table analysis as well as Pearson residuals were determined for “neuropathic pain”, “migraine with aura” and a chord plot with all genes implicated in the queried disease with PPI confidence scores (StringDB) as edge weights and the main differentiation stage affiliation as arc color was constructed. Standardized Pearson residuals were used to determine potentially enriched/depleted disease-associated genes for each differentiation stage. Human microRNA disease database (HmDD) was used to detect miRNAs associated with migraine disorders, chronic pain and neuropathic pain and temporal trajectories of associated miRNAs were determined and clustered using hierarchical clustering of z-scored vst counts and visualized as a heatmap along the time.

Since a number of severe human pain disorders are strongly associated with gain or loss-of-function mutations of ion channels in neurons (59–63), we applied the NOCICEPTRA tool to exemplarily investigate possible susceptibility windows during iPSC differentiation using the custom-gene set function for the two disease entities “*neuropathic pain*” and “*migraine with Aura”* evaluating the risk association score (association score >= 0.2) obtained from the OpenTarget.org database. The overall number of genes within each super-cluster was used as background to identify disease linked genes, and standardized Pearson residuals were calculated to determine the contribution of each cell to the contingency table statistics (**Figure 7B, Supplementary Table S15**). Supercluster enrichments for neuropathic pain and migraine with aura associated genes revealed a significant enrichment during the *nociceptor stage* for migraine with Aura, while for neuropathic pain the largest number of genes were associated with the nociceptor stage as expected, such as *ORPM1*, *CHRNA4*, *CACNA1B* and the metabotropic glutamate receptors *GRM5*, however no significant enrichment was detected (**Figure 7B,** contingency table statistics > 0.05; **Supplementary Table S19/20**).

Interestingly, several genes were abundantly expressed already during iDN differentiation and associated with Migraine with Aura such as *HTT*, *TOR1A* but also *DRD2* which suggests a putative implication in the development of neural progenitor cells (**Figure 7B**). Additionally, ion-channels associated with migraine with aura but also neuropathic pain were abundantly expressed in the nociceptor stage such as *SCN1A*, *SCN2A*, *SCN3A*, *SCN9A*, *KCNA1*, *HCN2*, *CLCN3* and *KCNA2* but also serotonergic receptors such as *HTRA2* and glutamatergic receptors such as *GRM5* and *GRIN3A*, proving suitability of the model to study both diseases (**Figure 7B, Supplementary Table S20**).

Mapping miRNAs associated with neuropathic pain, chronic pain or migraine to diseases using the human microRNA Disease Database (HMDD v.3.2, www.cuilab.cn) revealed two groups of miRNA which were associated with disease entities (**Figure 7B, righ panel)**. These were significantly expressed in early differentiation (D0-D5) and in nociceptor stage (D16-36). microRNAs associated with early differentiation putatively imply a high regulatory control of iDN in the differentiation and maturation. Interestingly, hsa-miR-155-5p was significantly associated with both disease entities “neuropathic pain” and “migraine” and further was associated with neurite outgrowth related genes in this study. We identified common targets associated with migraine and the hsa-miR-155-5p high confidence target space such as *PLP1*, *TRAK1,* SYNJ1 and *CREB5* (**Figure 7B**).

Based on these findings, it can be anticipated that impairments of migraine associated genes and associated miRNA regulation can be assessed experimentally already during early differentiation/neural progenitor phases of iPSC derived nociceptor models. The same appears to apply for genes associated with neuropathic pain questioning the role of those genes not only in mature nociceptor function but also nociceptor development and potential genetic impairments affecting both.

Hence, NOCICEPTRA was queried and compared against CORTECON expression patterns of important developmental marker genes during cortical neurogenesis which revealed distinct similarities and expected differences. As a first example, FGF receptors (FGFR1, FGFR2 and FGFR3) contribute significantly to corticogenesis and peak around the early corticogenesis in the CORTECON dataset (23), However, expression was only conserved for FGFR2 and FGFR3 in iDNs peaking around D5 which expectedly indicate neural crest cell generation common to both neuron types. By contrast, FGFR1 trajectories steadily decreased, suggesting that FGFR1 depression is necessary for nociceptor genesis only (**Supplementary Figure S10 A**). Second, WNT signaling is crucial for corticogenesis/cortex formation and WNT7B and WNT8A expression increases during cortical specification (23). This was not the case for iDNs where expression of WNT7B and WNT7A genes was low at all stages. Querying all WNT genes to elucidate potential susceptibility windows revealed that the majority of WNT genes (*WNT1*, *WNT4*, *WNT10B*, *WNT3A* and *WNT5B*) appeared to contribute to neural tube formation and neural crest cell development around D5 and D9 and that especially WNT1 is abundantly expressed (**Supplementary Figure S10 B,C**) (64, 65). The complex NOCICEPTRA and CORTECON data sets and tools can therefore be exploited for mechanistic studies dissecting differential developmental regulation of neuron subtypes in the peripheral and central nervous system.

## Discussion

Fueled by the urgent need for better and accessible human-based model systems, increasing efforts have been made to understand principal transcriptomic signatures responsible for the development of various tissue types, including cortical neurons by developing iPSC-derived differentiation protocols (23, 66). The majority of studies to date focus primarily on expression profiles of protein coding genes thereby disregarding the critical importance and application potential of miRNAs for the regulation of cell development and differentiation. In order to fill this gap, we performed the first unbiased temporal transcriptomic analysis of paired mRNA::miRNA expression during iPSC-derived nociceptor (iDN) differentiation in three different iPSC cell lines and detected several entities where miRNAs controlling an entire set of target genes apparently serve critical roles as regulatory master switches.

### mRNA and miRNA signatures characterize five distinct stages of nociceptor differentiation

Different iPSC donor lines and clones can react to differentiation protocols in a highly variable manner thereby impeding the identification and validation of conserved developmental pathways and molecular disease signatures (67) (16, 24). All three iPSC lines in our study developed into a nociceptor like phenotype and were functionally capable of firing action potentials with largely conserved transcriptomic profiles and highly correlative inter-module expression patterns. Expression trajectories with an increased expression of *RUNX1,* a reduction in TrkA expression but an increased expression of TAC1 were consistent with previous literature, and the applied protocol predominantly generated peptidergic sensory neurons (3, 19). Thereby, five independent stages of sensory neuron differentiation were characterized by representative marker genes such as *POU5F1* (pluripotency stage), *NEUROG1* (early differentiation stage), *NTRK1* (early-differentiation – neural progenitor stage), *SOX10* (neural progenitor stage) and *CLCN3/SCN9A* (nociceptor stage), as well as gene pathway enrichments for each stage (68–70). Characteristic trajectories of mRNAs and miRNAs emerged for the three relevant features of maturating nociceptors: *neurite growth*, *synapse* and *ion channel expression*.

### Hub genes and miRNAs orchestrating differentiation and fate determination

miRNA-driven posttranscriptional gene regulation is pivotal during neurogenesis and loss of the miRNA-synthesizing enzyme Dicer leads to impaired neural stem cell differentiation and abnormal brain development (71, 72). Several miRNAs and their target genes are well accepted regulators of neuronal development and differentiation at defined stages of nociceptor differentiation, such as members of the miR-302, the miR-125b and the let-7 families (73). In particular, the miR-17∼92 cluster, was highly expressed and associated with the early differentiation of iPSC-derived nociceptors (D5-D9). The miRNAs hsa-miR-20a-5p, hsa-miR-92a, hsa-miR-19b and hsa-miR-18a were highly regulated and associated with glutamatergic synapse formation, neuronal cell body and endomembrane system organization pathway enrichments. The strong association of high confidence targets with neurite outgrowth and neurotransmitter systems, moves the suppression of miR-302 into focus as a critically important mechanism for neuronal development and fate decision (74–76). However, also miRNAs sparsely investigated in neuron development, such as miR-106a/b, miR-199b and miR-504, were abundantly expressed, and presumably involved in suppressing pluripotency related genes, thereby inducing and maintaining the neuronal phenotype.

### Neurite formation related pathways controlled by hub genes and miRNAs

Neuronal differentiation is accompanied by significant morphological changes including development of filopodia and outgrowth of neurites/processes particularly in the neural progenitor stage (D9-D16). miRNAs appeared to be involved in certain aspects of neurite formation. The most abundantly expressed intragenic miRNAs hsa-miR-363-3p, hsa-miR-20b-5p, hsa-miR-106a-5p, which are all embedded in the lncRNA AC002407.2 (77), targeted a variety of pathways implicated in axon guidance and showed strong correlation with NEUROG1, supporting their importance for neuron morphology and neurite outgrowth (78, 79). Interestingly, miR-20b/miR-106 directly suppress *NEUROG2* (80), thereby indirectly favoring *NEUROG1* expression and nociceptor differentiation.

Half of all consensus modules associated with neuronal development, were implicated in neurite development and neurite growth. Within these, two classes of genes were overrepresented: DUSPs – dual phosphatases, and RABs – brain-associated Ras small G-proteins, which are well associated with neurite outgrowth (81–83). The antiproliferative miRNAs hsa-miR-31-5p but also hsa-miR-103a-3p were highly abundant expressed early in differentiation (84). Since they specifically target DUSP and RAB genes, these two miRNAs could act as main drivers balancing neurite growth in sensory neurons during nociceptor differentiation and maturation.

Neurite outgrowth is also regulated by Rho-related pathways, and *RHOC, RHOB* and *RHOJ* were highly correlated with CDK8, for which regulation of actin-cytoskeleton assembly is well established. This pathway appears to be unique for sensory neuron differentiation in humans: whereas *RHOA* is highly expressed up to postnatal day 1 in rodents, *RHOA* in human sensory neurons gradually declines, while *RHOC* increased in iPSC-derived nociceptors, suggesting opposing biphasic regulatory control of cytoskeletal assembly during differentiation (85, 86).

### miRNAs regulating ion channels in iPSCs and maturing iDNs

Our results further support the importance of the miR-17∼92 cluster, since it may act as a general ion-channel regulator by putatively downregulating an entire group of important ion channels, including the voltage-gated sodium channels SCN2A and SCN9A, the pacemaker channels HCN1 and HCN2, the voltage-gated potassium channels KCNA1, KCNB1 and KCNH5, the inward-rectifier KCNJ6 as well as TRPM4, most of which are dysregulated in neuropathic pain or implicated in neuronal development, Thereby putatively protecting for excitotoxcity (87, 88). In particular the target-space of miR-20a (a member of the miR-17∼92 cluster) included multiple genes specifically regulating neurite outgrowth, making it a critically important hub miRNA for nociceptor differentiation. However, we also detected new sets of miRNAs with a putative role in targeting ion channels. miR-182 and miR-26b targeted the gap-junction related channels GJC1 and GAJ1, while the miR-20b-3p (and the miR-17∼92 cluster) and the miR-302 family were found to target SCN9A, TRPM7, or CLCN5-CLCN7 channels. GJC1 and GJA1 might be of particular importance for retaining pluripotency as they facilitate the reprogramming efficiency towards iPSCs (89). Also, the expressed TRPM7 channel appeared to be strongly controlled by at least 17 miRNAs during differentiation and this stresses its role in neuronal differentiation in particular of mechanosensitive sensory neurons through however unknown mechanisms (90, 91).

At the more mature stages, several ion channels emerged as targets of miRNAs, whose relevance for the transduction of painful stimuli, excitability or action potential propagation is well accepted. Two important nociceptor ion channels *SCN2A* and *SCN9A* (57, 58) were suppressed in immature stages by highly expressed miRNAs of the miR-17∼92 cluster which decreased during differentiation, while *SCN2A* and *SCN9A* expression increased. Also, the chloride transporters and channels *CLCN4-6* were strongly targeted by miRNAs especially by the miR-302 family. In particular the miR-302 family suppressed 10 ion-channel targets. This moves the miR-302 family into focus as a critical and efficient hub regulator mechanism suppressing entire clusters of channels and other machineries that are silenced in pluripotent cells.

### Differentiation of the synaptic release machinery

Although native primary sensory afferents do not form autapses (92), iDNs express synaptic proteins (18). This is surprising but may be related to the unique feature of nociceptors to release neuropeptides and glutamate from differing vesicle pools at peripheral and spinal terminals (93). Two paralogs of the calcium-dependent activator protein for secretion are priming factors for synaptic vesicles containing glutamate and large dense-core vesicles containing neuropeptide. Essential components of the synaptic release machineries such as syntaxin, synapsin, SNAP91 as well as different RAB proteins were subject to the control by hub miRNAs. While hsa-miR-302a-5p is essential for neuronal differentiation and neural tube closure the present study for the first time hints toward the implication of the miR-25 family in nociceptor development and suppression of synapse machinery related genes (94–97). Yet, it is unclear how the exocytotic release machinery from these two vesicle types in nociceptive neurons is differentially regulated, wherein synaptic transmission and neuropeptide release are equally important (98). To further investigate timepoint specific mRNA and miRNA regulation, we developed the open access NOCICEPTRA tool to extract expression patterns and time trajectories of miRNAs and their target gene space for mechanistic studies.

### Querying gene trajectories using the NOCICEPTRA tool

Based on disease enrichment data, we found that a set of genes and miRNAs contributing to nociceptor development was also associated with chronic pain and migraine. To further extend the current knowledge on disease-relevant gene and miRNA trajectories during nociceptor differentiation, we developed the NOCICEPTRA tool to visualize stage specific gene enrichments for diseases and KEGG pathways, as well as for single miRNAs. The NOCICEPTRA resource allows to explore pain and other sensory neuron-related disorders through the discovery of disease onset and susceptibility windows. The NOCICEPTRA platform is provided as a containerized online tool that enables the scientific community to retrieve the entire dataset of the experimentally determined expression trajectories, as well as to query custom gene sets of interest for pathway and disease enrichments. Querying our resource to differentiate expression patterns of important developmental marker genes identified during cortical neurogenesis (CORTECON) revealed distinct similarities and dissimilarities. As an example, the FGF receptor FGFR1 significantly contributed to corticogenesis but not nociceptor development, while FGRF2 and FRGR3 were found important for both nociceptor genesis and corticogenesis. This pinpoints the added value of CORTECON and NOCICEPTRA tools to generate novel mechanistic insight into neuron subtype programming.

Together, our findings suggest that miRNA::mRNA interactions act as critically important hubs for suppressing mature protein coding gene sets during pluripotency and for silencing pluripotency genes when neurons serve their mature function in the nervous system. This posttranscriptional regulation via miRNAs emerges as effective and of particular importance for phenotype decisions, neurite growth and target finding as well as synaptic processes for nociceptor interaction with target tissues and second order neurons. In order to fathom the complexity of the modules involved in these unique functions we provide an open resource for the scientific community that can be used to query pathways and miRNA-controlled gene sets.

### Conclusion

In summary our paired RNA and small RNA sequencing approach allowed us the complex mapping of a developmental gene and miRNA expression atlas discovering major hub genes and miRNAs significantly contributing to the development of iDNs. Thereby miRNAs and hub genes with specific roles in synapse development, neurite outgrowth as well as ion-channel development were identified and the resource NOCICEPTRA generated allowing general exploration of gene and miRNA expression, hub modules and pathways as well as disease susceptibility windows.

## Funding

This study was funded by the Austrian Science Fund FWF (P30809 to KK; DK-SPIN, B16-06 to MK) and the European Commission (ncRNAPain, GA 602133 to MK).

## Acknowledgement

The authors thank C. Wild and M. Tschugg of the IT Services, Medical University of Innsbruck, for providing general support regarding the computing environment.

## Author contributions

MZ, CLS, TK, KK and GK performed the experiments; MZ and KK developed the NOCICEPTRA tool; MZ, KK and MK conceptualized the study; MZ, KK, ZC and MK wrote the manuscript.

## Competing Interest

The authors declare no competing interests.

## Materials and Methods

### Induced pluripotent stem cell (iPSC) lines

Three different iPSC lines, adult clone 2 (AD2; SFC-AD2-1, male), adult clone 3 (AD3; SFC-AD3-1, female) and clone 840 (SFC-840-03-01, female), derived from healthy donors with no known history of neuronal diseases as we described (18), were used throughout the experiments for reprogramming, and paired RNA and small RNA co-sequencing (**Supplementary Table S16**). Cells were stored in liquid nitrogen until thawing. After thawing cells were passaged twice and maintained in mTeSR1 medium at 37°C and 5% CO2.

### iPSC-derived sensory neuron (iDN) differentiation

AD2, AD3 and 840 iPSCs were cultured on Matrigel (Corning) coated 6-well plates and maintained in mTeSR1 medium (STEMCELL Technologies). Nociceptor differentiation was performed as previously published (18, 19). In brief, approximately 80 x 10^4^ cells per 6-well plate were seeded as single-cells at passage 2 after thawing. Cells were maintained overnight in mTeSR1 medium containing the Rho-associated kinase (ROCK) inhibitor Y-27632 (Sigma-Aldrich) until cells reached 60-70 % of confluency. Subsequently, dual-SMAD inhibition was initiated by inhibiting BMP4 and TGF-β in knockout serum replacement (KSR; Knockout DMEM [Gibco], 15% KO serum replacement [Gibco], 1% non-essential amino acids (PAA), 2mM Glutamax [Gibco], 1x Penicillin/Streptomycin [Sigma-Aldrich], and 100µM !-mercaptoethanol [Gibco]) medium supplemented with 100 nM of LDN-1931189 (STEMGENT) and 10 µM SB431542 (selleckchem.com) for 5 days. Additionally, 3 µM CHIR99021 (GSK3beta inhibitor, selleckchem.com), 10 µM DAPT (gamma-secretase inhibitor, selleckchem.com) and 10 µM SU5402 (VEGFR-2, FGFR-1 and PDGFRB inhibitor, selleckchem.com) were added from day 2 (D2) to D12. From D4 on N2/B27 (Neurobasal medium^TM^/GLUTAMAX/ 2% B27 w/o vitamine A [Gibco]/ 1% N2 [Gibco]) was added to the KSR medium every second day in increments of 25%. On D12, 80 x 10^4^ cells were passaged onto Matrigel coated 6-well plates according to the supplier’s recommendations and maintained in N2/B27 medium supplemented with 25 ng/ml nerve growth factor (hNGF, PreproTech) 25ng/ml brain derived neurotrophic factor (hBDNF, PreproTech) and 10 ng/ml glia derived neurotrophic factor (hGDNF, PreproTech). Y-27632 was added for one day following passaging, to promote survival of neuronal-like progenitor cells. Cytosine-β-D-arabinofuranoside (4 µM, Sigma Aldrich) was administered on D14 for ∼24 h to reduce the amount of non-differentiated proliferating cells. Cultures were maintained until D36 and medium was changed every other day.

### Time-course paired RNA and small RNA sequencing

We performed paired RNA and small RNA sequencing to identify temporal changes of gene expression during the development of nociceptor differentiation. Data were generated for six different time-points (i.e., D0, D5, D9, D16, D26 and D36) chosen based on morphological rearrangement such as major treatment changes during the protocol and neurite outgrowth in all three different iPSC cell lines (i.e., AD2, AD3 and 840; N=18 per cell line).

To explore gene and miRNA expression patterns during the development of iPSC-derived nociceptors, cells were collected in triplicates for each cell line. To harvest cells, phosphate buffered saline (PBS) was used to wash off the remaining medium and to reduce the amount of cell debris. Cells were detached with 1x Tryp-LE Express (Sigma) for 3 minutes. Tryp-LE was inactivated with PBS in a 1:1 mixture and cells were centrifuged at 500 g for 5 minutes. Subsequently, supernatant was depleted and cells were snap frozen in liquid nitrogen and stored at -80° C until further processing. RNA and small RNA co-sequencing was performed by IMGM Laboratories (Martinsried, Germany). To avoid batch effects, all RNA and small RNA samples were processed in paralell. Briefly, in order to isolate RNA and small RNA from frozen samples, the miRNeasy Mini Kit (QIAGEN) was used according to the manufacturer’s instructions. RNA concentration and purity were assessed using NanoDrop ND-1000 spectral photometer (Peqlab) and the quality of the RNA was determined using RNA 6000 Nano LabChip Kits (Agilent Technologies) on a 2100 Bioanalyzer (Agilent Technologies). RNA quality based on RNA integrity (RIN) is presented in **Supplementary Table S17**. Furthermore, all libraries were quantified using the highly sensitive fluorescent dye-based Qubit ds DNA Assay Kit (Qiagen). Whole RNA library preparation was performed using the TruSeq stranded mRNA HT technology (Illumina) according to the manufacturer’s protocol. All single mRNA libraries were pooled into one sequencing library pool with an equal amount of DNA per sample. After quantification, the final sequencing library was diluted to 2.25 nM followed by denaturation with NaOH. Small RNA sequencing was performed using the NEBNext small RNA Library Prep Kit for Illumina using 500 ng total RNA, according to the manufacturer’s instructions. Purified small RNA sequencing libraries were quantified using the fluorescent dye-based Qubit ds DNA HS Assay Kit (Thermo Fisher Scientific) on a 2100 Bioanalyzer. After quantification, 2 nM of the sequencing library were used for further analysis. Both RNA and small RNA libraries were clustered and sequenced using the Illumina NovaSeq600 and performed using single reads with a length of 75 bp. Technical quality parameters were evaluated using the SAV software and revealed that more than 85% of bases passed the Q30 bases threshold for all samples.

### RNA read processing and differential gene expression analysis

Derived mRNA FastQ files were aligned to the human reference genome (GRCh38.p13, release 33) provided by GENCODE (www.gencodegenes.org) using *STAR* (v2.7) software with the default parameters (99). Small RNA sequencing libraries were prepared using the NEBNext Small RNA Library Prep Set for Illumina. To further process small RNA reads adapter sequences (AGATCGGAAGAGCACACGTCTGAACTCCAGTCAC) were trimmed using *Flexbar* v3 (100) and aligned to the human reference genome (Parameters can be found in **Supplementary Table S18**). Raw counts for mRNA and small RNA were obtained sorted *.bam* files using *HTSeq* (v 0.11.1). Raw mRNA and miRNA counts were determined by annotation to the GENCODE comprehensive gene annotation file (GRCh38.p13_chr_patch_hapl_scaf.annotations.gtf, www.genecodegenes.org) and the *hsa.gff* file provided by miRbase (release 22.1, www.mirbase.org), respectively.

### Differential gene expression

Detection of differentially expressed genes (DEG) and miRNAs (DEmiR) was performed using the Bioconductor R-package *DESeq2* (release 3.10) and RStudio (101). A first model was deployed merging all samples from each timepoint and each cell line, and evaluating differential gene and miRNA expression using the likelihood ratio test (LRT) with timepoint as independent parameter, to detect significant changes at any timepoint during the differentiation. A second model was deployed determining the number of DEGs and DEmiRs for each cell line at any given timepoint. Quantification of DEGs and DEmiRs, as well as log2-fold changes were estimated using the normal shrinkage function, and the FDR-corrected significance threshold was set to FDR < 0.05 for DEGs and FDR < 0.1 for DEmiRs. Normalized counts were further variance stabilized using the vst function.

To identify samples with an increased similarity of gene and miRNA expression, principal component analysis (PCA) as well as hierarchical clustering were applied on log-transformed counts. To determine the number of DEGs for each gene biotype, the Bioconductor R-package *BioMart* was used (102) to assign the biotype to each gene.

### Weighted gene co-expression network analysis (WGCNA)

A weighted gene co-expression network analysis (WGCNA) was used to detect modules of genes that are highly correlated (103). To be as unbiased and unsupervised as possible, normalized read counts obtained from the DESeq2 analysis for all mRNAs and miRNAs were filtered for genes and miRNAs that had at least 5 counts in 3 samples using Python’s “Numpy” and “Pandas” package. A consensus dataset for genes and miRNAs was constructed separately using an inner joint on shared filtered genes and miRNAs for all three cell lines and quality as well as consistency of the dataset was quantified using the *checkSets* function as well as the *goodSamplesGenesMS* function integrated in the WGCNA package. Sample trees for each of the three cell lines were constructed using the *hclust* function. The *blockwiseConsensusModules* function (softthresholding power = 16 (mRNAs)/7 (miRNAs), maxBlockSize = 37000, network-Type = “signed”, minModuleSize = 30, mergeCutHeight = 0.15, minKMEtoStay = 0.2 (mRNA)/0.0 (miRNA)) was used to construct modules of significantly correlated genes and miRNAs. Modules of gene expression were defined as a cluster of densely interconnected genes based on co-expression. The co-expression matrix was determined by Pearson correlation between two genes and transformed into an adjacency matrix, using a soft threshold determined according to the scale-free topology criterion using the following formula:

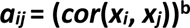

where *cor(x_i_,x_j_)* is the correlation of any possible gene pair and *b* ist the soft-threshold power to generate a weighted network adjacency matrix.

This matrix was used to determine the topological overlap metric (TOM), which was then further used to cluster genes and miRNAs according to their expression trajectories.

The multiSetMEs function was used to obtain metrics about the correlation and the preservation of the consensus modules across the three cell lines and the inter-module correlation within each cell line. Hub genes were identified extracting the *kModule Eigengene* for each gene in each module. The grey module, representing genes which could not be assigned to a specific module and was therefore excluded from further analysis.

### Supercluster analysis and Singular Value Decomposition

Identification of Eigengene modules generated a high number of modules (49 modules -> mRNA, 17 modules -> miRNA), of which subsets exhibited similar gene-expression patterns. To collect modules with correlated gene expressions, hierarchical clustering using the *“average”* agglomerative clustering algorithm (mRNA distance: 3.2, miRNA distance 3) was used on module-wide averaged z-scored standardized variance stabilized (vst) gene expression counts, which were further averaged across the three cell lines for genes and miRNAs. Furthermore, a second sanity check approach was employed using singular value decomposition on z-scored standardized vst gene expression values averaged across the cell lines, according to Bennett et al. (104) using the python framework “scikit learn” by decomposing the gene and miRNA expression matrices.

### Gene pathway enrichment and disease enrichments

Every module and each supercluster underwent gene pathway enrichment analysis as well as disease enrichment analysis (DOSE, DisGeNet, www.disgenet.org). g:Profiler (https://biit.cs.ut.ee/gprofiler/gost) was used for the 4 annotation spaces Gene ontology (GO) biological process (GO:BP), GO cellular components (GO:CC), GO molecular function (GO:MF) and Kyoto Encyclopedia of Genes and Genomes (KEGG) by reducing the number of significantly enriched pathways to best per parent by a custom written script, as described previously (77, 105). To further reduce the complexity of gene pathway enrichments, only the top 5 GO:BP, GO:MF, GO:CC and KEGG pathways were visualized using the negative log10(p-value).

The Bioconductor R-package *DOSE* was used to determine potential disease enrichments for each supercluster/module (parameters: p-value cutoff: 0.01, pAdjustMethod: “BH”, minGSize = 10, maxGSSize = 10000, qvalueCutoff = 0.2) and the *DisGeNET* database (www.disgenet.org) was used to determine miRNA specific disease enrichments (106, 107), using contingency analysis provided by the python framework “statsmodel” and the function contingency table. Enrichments and contributions of each individual differentiation stage were determined by means of standardized Pearson Residuals using “statsmodel” and a Pearson residual above > abs(+-2) were considered as putative stage specific enrichment or depletion.

### Reversed miRNA targeting prediction

To evaluate potential miRNA targets implicated in the differentiation of iPSC-derived nociceptors, each mRNA::miRNA time trajectory (z-score standardized vst counts) was correlated using *Pearson correlation*. Additionally, the databases StarBase (108), miRTarBase (109) and TarBase (110) were used to determine the validation status, and DIANAs microT-CDS algorithm (111) was used to predict a potential interaction. Moreover, we ranked miRNAs in percentiles based on their baseMean expression during the differentiation, to take the amount of available miRNA molecules for targeting into consideration. All the described parameters were used to determine an overall target score, calculated with the following formula:

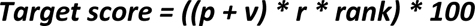

where *p* is the microT-CDS v5 score considered only if score > 0.9, *v* is a boolean variable where 1 is validated and 0 is not validated yet, *r* is the Pearson correlation coefficient between the miRNA and the gene and *rank* is the percentile rank of the miRNA based on its expression (77).

### miRNA interaction networks

miRNA:mRNA interaction network analysis, for miRNAs known to play a prominent role in the regulation of iPSC-derived nociceptor differentiation, was performed for each expressed miRNA (BaseMean > 5 counts in at least 3 samples).

Only genes::miRNA interactions that were highly negatively correlated (r < -0.7) were taken into consideration. Those genes that were at least validated or predicted (DIANA microT-score > 0.9) with an overall target score > 50 if not differently stated were used for further stage enrichment analysis. Target gene PPI were identified using StringDB with the confidence score set to 0.7 if not stated otherwise and a chord plot with genes as nodes, the arc color as supercluster affiliation and the edges as PPI was constructed using “Holoviews”, “Matplotlib” and “Bokeh”. To identify potential supercluster stage enrichments the number of targeted genes for each supercluster was determined and compared to the global number of possible targeted genes per differentiation stage. A contingency table was constructed and a χ^2^ test performed. To evaluate which modules were potentially enriched, standardized *Pearson Residuals* were determined and a *Pearson Residual* cutoff > 2 was used to distinguish highly enriched differentiation/module stages. To identify the regulation of functional pathways by single miRNAs g:Profiler analysis of the targeted genes were performed.

For ion-channel specific miRNA::mRNA interactions a spring-forced network was constructed using the target-score as edges weight. Networks were drawn with the python module “NetworkX” and refined with Cytoscape and the NetworkAnalyzer.

### Hub genes:hub miRNAs interactions

Genes considered hub genes are characterized by the k-module eigenvector (kME) of above > 0.8 and a p-value < 0.05. Hub-gene networks for each module were constructed using the top 30 hub-genes ranked based on the kME and the p-value for all three cell lines and the edges weight was defined as the correlation between two genes calculated using “scikit learn” stats.pearsonr() function.

Only edges with a correlation > 0.8 were considered and networks were constructed using *NetworkX using the kamada kawai layout*. Hub miRNAs are miRNAs which target hub-genes. To determine hub genes that share the same miRNA interaction space, an adjacency matrix using the target score (> 70) for each miRNA::mRNA interaction was constructed and dimensional reduction along the gene axis was performed using the python module unifold manifold approximation and projection “UMAP” (n_neighbor = 20, min_dist = 0.1, metric = “manhattan”). To cluster genes potentially targeted by the same set of miRNA, kMeans clustering (n_cluster = 8) was performed using “scikitlearn” kmeans() function. We further performed g:Profiler enrichment analysis to determine k-Means cluster enriched for neuronal pathways.

### Indirect immune fluorescence microscopy

AD3 (SbAd-03-01) and 840 (SFC840-03-01) iPSC clones differentiating into iDNs were fixed on D0, D16, D26, D36 with 4% paraformaldehyde in PBS at room temperature, for 10 min. Fixed iDNs were blocked with 5% normal goat serum (NGS) in PBS supplemented with 0.2% BSA and 0.2% Triton X-100, for 30 min. Primary antibodies were applied at 4 °C overnight in a humidified chamber, and detected by fluorochrome-conjugated secondary antibodies (Alexa goat anti-rabbit A594 (#A32740), goat anti-mouse A594 (#A32742), goat anti-rabbit A488 (#A32731), goat anti-mouse A488 (#A32723), and goat anti-rat (#A11007, all 1:4000; Thermo Fisher Scientific). Primary antibodies used were anti-TUJ1 (1:600, mouse monoclonal, R&D Systems, #MAB1195); anti-Substance P (SP) (1:1000, mouse monoclonal Abcam #SP-DE4-21), anti-CGRP #149598 (1:500, rabbit polyclonal, Cell Signalling), anti-Nav1.7 (SCN9A) #ASC-008 (1:1000,rabbit polyclonal, Alomone Labs), anti-PIEZO2 #APC-090 (1:500, rabbit polyclonal, Alomone Labs), anti-panTRK #ab76291,(1:500, rabbit monoclonal, Abcam), anti-NANOG #ab62734 (1:500, mouse monoclonal, Abcam), anti-Ki-67, #ab156956,(1:250, rat monoclonal, Abcam), anti-TMEM119, #ab209064, (1:500, rabbit monoclonal, Abcam), anti-PERIPHERIN, #ab4666, (1:500, rabbit polyclonal, Abcam), anti-GFAP #04-1062 (1:250, rabbit monoclonal, Merck Millipore). Nuclei were counterstained with DAPI (4ʹ,6-diamidino-2-phenylindol) 1:10,000 (Thermo Fisher Scientific). Images were recorded using an Axioimager 2 Microscope with cooled CCD camera (SPOT Imaging Solutions) for glass coverslips and an inverted Axiovert

200 M setup for Ibidi dishes (both Carl Zeiss Light Microscopy). Average fluorescence intensities were quantified using Metaview software (version 7.8.0.0, Molecular Devices, LLC, San Jose, CA95134) with the line scan plug-in to quantify fluorescent intensities along a defined line cutting individual neuronal structures. Cell counts positive and negative for primary antibodies were correlated to DAPI positive nuclei and compared between the individual differentiation days. In general 2 dishes/cellline/condition (Day 16-D 36) were stained and analyzed in at least 2 individual

### RT-qPCR analysis

Reverse transcription and qPCR reactions were performed on the same samples that were used for sequencing according to the protocol provided by the supplier (Thermofisher scientific). In short, each reverse transcription reaction contained 10 ng of total RNA, 1X reverse transcription buffer, 5.5 mM MgCl2 (GeneAmp 10X PCR Buffer II & MgCl2, #N8080130), 1 mM dNTPs, RNase inhibitor (#N8080119), 50 units of MultiScribe Reverse Transcriptase (#4311235) and 1X RT specific primers (**Table 2**) volume of adjusted to 15 µl with nuclease free water (#R0582). The RT program was set to 30 min at 16°C, 30 min at 42°C, 5 min at 85°C, followed by a holding step at 4°C. After reverse transcription, reactions were stored overnight at -20°C. Each qPCR reaction contained 1.33 µL of the RT product, 1X TaqMan® Universal Master Mix II, no UNG (#44440049), 1X of the appropriate assay (**Table 2**) and nuclease free water up to a final volume of 20 µL. Reactions for each sample were prepared as technical duplicates, alongside reverse transcription non-template controls, loaded on MicroAmp Fast Optical 96-well reaction plates (#4346906) and placed in the QuantStudio 6 Pro Real-Time PCR system (all reagents for RT-qPCR were purchased from ThermoFisher Scientific). The PCR cycle protocol was: 10 min at 95°C, 40 two-step cycles of 15 s at 95°C and 1 min at 60°C. In order to account for potential methodological variabilities all RT (day 1) and qPCR reactions (day 2) were processed in parallel. Threshold was set at 0.1 and results were obtained from the QuantStudio Design and Analysis software. The expression levels of the miRNA of interest were quantified and expressed as relative expression to the respective expression of the reference gene (U6) using the 2^-ΔCq^ method.

**Table 1:** Protein-protein interaction network of each module.

module
| **module** | **nodes** | **PPI enrichment p-value** |
| --- | --- | --- |
| *bisque 4* | 58 | 3.60E-03 |
| *black* | 584 | 2.23E-11 |
| *blue* | > 2000 | n.d |
| *brown* | > 2000 | n.d |
| *brown4* | 75 | 5.92E-02 |
| *cyan* | 231 | 2.45E-08 |
| *darkgreen* | 75 | 1.00E-16 |
| *darkgrey* | 137 | 2.04E-06 |
| *darkmagenta* | 124 | 1.00E-16 |
| *darkolivegreen* | 114 | 1.21E-01 |
| *darkorange* | 121 | 3.47E-01 |
| *darkorange2* | 65 | 3.52E-01 |
| *darkred* | 179 | 3.20E-02 |
| *darkslateblue* | 41 | 7.13E-01 |
| *darkturquoise* | 145 | 3.75E-09 |
| *floralwhite* | 76 | 1.63E-05 |
| *green* | 813 | 1.00E-16 |
| *greenyellow* | 324 | 1.00E-16 |
| *grey60* | 217 | 1.45E-07 |
| *ivory* | 82 | 2.98E-04 |
| *lightcyan* | 232 | 1.23E-05 |
| *lightcyan1* | 78 | 1.00E-16 |
| *lightgreen* | 195 | 3.60E-05 |
| *lightsteelblue1* | 88 | 2.94E-05 |
| *lightyellow* | 184 | 5.80E-04 |
| *magenta* | 384 | 1.00E-16 |
| *mediumpurple3* | 88 | 2.58E-02 |
| *midnightblue* | 235 | 1.54E-01 |
| *orange* | 154 | 3.82E-05 |
| *orangered4* | 76 | 4.21E-14 |
| *paleturquoise* | 129 | 1.00E-16 |
| *pink* | 351 | 7.24E-05 |
| *plum1* | 95 | 1.94E-01 |
| *plum2* | 44 | 2.70E-01 |
| *purple* | 307 | 7.37E-08 |
| *red* | 785 | 1.00E-16 |
| *royalblue* | 170 | 3.26E-09 |
| *saddlebrown* | 139 | 3.14E-03 |
| *salmon* | 234 | 1.00E-16 |
| *sienna3* | 85 | 1.00E-16 |
| *skyblue* | 125 | 1.16E-07 |
| *skyblue3* | 107 | 1.64E-02 |
| *steelblue* | 133 | 1.75E-01 |
| *tan* | 258 | 1.00E-16 |
| *turquoise* | > 2000 | n.d |
| *violet* | 133 | 1.00E-16 |
| *white* | 152 | 1.09E-07 |
| *yellow* | >2000 | n.d. |
| *yellowgreen* | 92 | 1.01E-06 |

**Table 2:** miRNA TaqMan Assay IDs.

| miRNA | Assay ID |
| --- | --- |
| hsa-miR-25-3p | 000403 |
| hsa-miR-103a-3p | 000439 |
| hsa-miR-146a-5p | 000468 |
| hsa-miR-302a-5p | 002381 |
| hsa-miR-335-5p | 000546 |
| Reference gene |  |
| U6 | 001973 |

### Whole-cell patch clamp analysis

Single cell voltage-clamp as well as current-clamp recordings were performed using the whole cell configuration of the patch clamp technique. Cells were kept in extracellular solution (ECS) containing (in millimolar); NaCl (150), KCl (5), CaCl2 (2), MgCl2 (1), HEPES (10), Glucose (10), and the pH was set to 7.3 with NaOH. Borosilicate glass pipettes (Science Products) were pulled with a horizontal puller (P-97, Sutter Instruments Company) with an average pipette resistance of 5 MOhm after filling with intracellular solution (ICS) (in millimolar): K-Gluconate (98), KCl (50), CaCl2 (0.5), MgCl2 (2), EGTA (5), HEPES (10), MgATP (2), NaGTP (0.2), and pH adjusted to 7.3 with KOH. Serial resistance (R-series) was compensated in all recordings using the automatic compensation function provided by the HEKA Patchmaster (> 60%, 10µs). Recorded neurons were held at -60 mV for voltage-clamp recordings and 10 mV depolarizing voltage pulses applied from -60mV to 40mV to assess peak (minimal current value per step) as well as sustained current amplitudes (mean value per step). The current-clamp mode was used to retrieve the resting membrane potential at 0pA. The minimal current to induce an action potential (Rheobase) was obtained using increasing 5pA depolarizing pulses. Action potential parameters analysed were duration (from threshold to repolarization), repolarization, threshold onset and half-width using the integrated AP fitting function of the HEKA Fitmaster. All recordings were performed in AD3 and AD2 at D36 platted on Matrigel coated IBIDI dishes maintained in N2/B27 Media supplemented with NGF, BDNF and GDNF.

### Statistics

Statistical analysis was performed using the Python packages *sklearn*, *statsmodel*, *numpy*, *pandas*, *networkX and requests* as well as the Bioconductor R packages *DOSE* and *WGCNA*. Differential gene expression analysis was performed using a hierarchical model approach implemented in the Bioconductor package *DESeq2* with the threshold of significance after FDR correction set to FDR < 0.05 (mRNA) and FDR < 0.1 (miRNA).

WGCNA consensus module affiliation was calculated by means of the WGCNA package distributed by Bioconductor, using kME and metakME and the module associated p-value for a significant affiliation was set to < 0.05.

χ^2^ test was performed following contingency table analysis with the p-value threshold set < 0.05. Pearson residuals were calculated using Python’s “statsmodel”, and Pearson residuals above and below abs(2) were considered as indicator for enrichments/depletions. Correction for multiple comparisons was performed where appropriate using the non-negative *Benjamini-Hochberg* FDR correction.

### Key Resource Table

A key resource table as well as software used for the project is deposited in **supplementary Table S21.**

### Availability of Data and Materials

Raw RNA and small RNA sequencing datasets are made available at Gene Expression Omnibus database (GEO; accession number GSE161530). All tables as well as the source code for the DESeq2 analysis and the WGCNA analysis are available in the supplement, and the project-specific GitHub page (https://github.com/MUIphysiologie/NOCICEPTRA). StarBase, miR-TarBase and TarBase databases necessary for reanalysis of the work can be obtained on the respective webpages, or on request.

Additionally a containerized tool will be available for further analysis (https://hub.docker.com/repository/docker/muiphysiologie/nociceptra_mui) using Python 3.8.2 and Streamlit (v.0.70, www.streamlit.org), a tool for rapid front-end development for data-science related projects. The tool can be easily accessed using Docker for Desktop for Windows, Linux and MacOS. The miRNA-Edges database can be provided on request or downloaded via the NOCICEPTRA docker tool (Download section), since size of the database is exceeding GitHub repositories limits.

## Supplementary Figure Legend

**Supplementary Figure 1: (A)** Hierarchical clustering of gene expression of each individual cell line per timepoint. **(B)** Intersection analysis of differential expressed genes are visualized in the Venn Diagram for AD2, AD3 and 840 clones. **(C)** WGCNA analysis report, with scale free topology and soft threshold parameters as well as mean and median connectivity

**(D)** Expression trajectories of the neuropeptides (TAC1, CALCA and CALCB)**. (E)** Immunostaining of CGRP (CALCA, red), peripherin (PER, green), DAPI and the respective linescan-scatterplot at D36 indicating colocalization of CGRP and PER in the cytoplasm and neurites. **(F,G)** Literature described murine nociceptor and developmental marker trajectories in iDNs. **(H)** Temporal trajectories of human DRG enriched transcription factors as described in Ray et al. (2018). **(I)** Spearman correlation analysis of log2 TPMs for iDNs and human cultured DRG (hDIV) and native human DRG (hDRG) obtained from Wangzhou et al. (2020). **(J,K)** Temporal trajectories of cultured human DRG enriched kinases and g-protein coupled receptors, illustrated by means of TPM.

**Supplementary Figure 2: (A)** Triple label immuofluorescence stainings of SCN7A (Nav1.7), TAC1 (Substance-P), NTRK1-3 (panTRK) and PIEZO2 costained with peripherin, or Tuj1 and DAPI at D16, D26 and D36 during iDN differentiation in AD3 and 840 cells. **(B)** Fluorescence intensity of the markers was measured within the respective field of view and compared between the timepoints. **(C)** Distribution analysis of positively and negatively stained cells within the field of view and comparison between the individual differentiation days. iDN differentiations.

**Supplementary Figure S3 (A)** Triple labeling immunofluorescence staining using either NANOG or KI-67 together with peripherin (PER) and DAPI. Arrowheads in the upper pannel indicate positively stained NANOG/KI-67 cells at D0. Arrowheads in the lower panel indicate NANOG/KI-67 negative cell cohorts at D36. IF positive cells for NANOG and KI-67 were compared to all DAPI positive cells at D0 and D36 and stacked barplots generated.

**(B)** Triple labeling of TMEM119 or GFAP together with TUJ1 and DAPI at D36 of iDN differentiation. **(C)** Temporal trajectories of GFAP and TMEM199 indicated low (TMEM119) and absent expression of GFAP

**Supplementary Figure 4: (A)** Current-voltage relationship accessed for two celllines (AD2, AD3, n = 15 cells) by means of a step-protocol (-60mV – 40mV) and peak as well as sustained current calculated and normalized against the membrance capacitance. **(B)** Resting membrane potential was evaluated in current clamp mode with a 0 pA injected current over 1 minute and three intervals (start (mean_1), middle (mean_2), end (mean_3) selected. **(C)** All cells functionally accessed, were capable of firing action potentials and a representative figure made (n = 10 (AD3)/ 11 (AD2) cells). **(D)** Action potentials were fitted by means of the integrated fitting function implemented in HEKA Fitmaster and Action Potential Duration, Onset as well as duration from onset till repolarization (mindT) investigated. **(H)** Differential expression analysis of voltage-gated sodium channels at D36 between AD3 and AD2

**Supplementary Figure 5 - Gene Trajectories:** Z-scored standardized gene expression trajectories for each individual module, for each cell line (blue: 840, red: AD2, green: AD3, respectively). X-axis represent differentiation dates, y-axis z-standardized vst-counts

**Supplementary Figure 6 - Gene Trajectories:** Z-scored standardized gene expression trajectories for each individual module, for each cell line (blue: 840, red: AD2, green: AD3, respectively). X-axis represent differentiation dates, y-axis z-standardized vst-counts.

**Supplementary Figure 7 - Gene Trajectories:** Z-scored standardized gene expression trajectories for each individual module, for each cell line (blue: 840, red: AD2, green: AD3, respectively). X-axis represent differentiation dates, y-axis z-standardized vst-counts.

**Supplementary Figure 8 (A,B)** PCA and hierarchical clustering of miRNA expression for each individual cell line. **(C)** Intersection analysis of differential expressed miRNAs are visualized in the Venn Diagram for AD2, AD3 and 840 clones. **(D)** WGCNA parameter report for soft threshold power, median and mean connectivity and max connectivity. Soft threshold topology model fit reaching 0.9 is achieved after the soft-threshold was set to 7. **(E)** WGCNA module preservation reports revealed high preservation across most modules detected with an average preservation score of 0.87. **(G)** qPCR validation of 5 miRNAs (n = 3/timepoint/cellline) enriched at distinct differentiation stages.

**Supplementary Figure 9 - miRNA curves:** Z-scored standardized miRNA expression trajectories for each individual WGCNA miRNA module for each cell line (blue: 840, red: AD2, green AD3).

**Supplementary Figure S10 (A)** Temporal Trajectories of FGFR1-3 using vst standardized counts and tpm indicative of relative expression. **(B)** Temporal expression of WNT related genes using vst counts and tpm indicative of relative expression. **(C)** Temporal expression of WNT related genes highly expressed around D9 using vst counts and tpm indicative of relative expression

## Notes

https://hub.docker.com/repository/docker/muiphysiologie/nociceptra_mui

https://github.com/MUIphysiologie/NOCICEPTRA

### Competing Interest Statement

The authors have declared no competing interest.

