## Supplementary figures and images for "NOCICEPTRA: Gene and microRNA signatures and their trajectories characterizing human iPSC-derived nociceptor maturation"

### Supplementary Figure 1

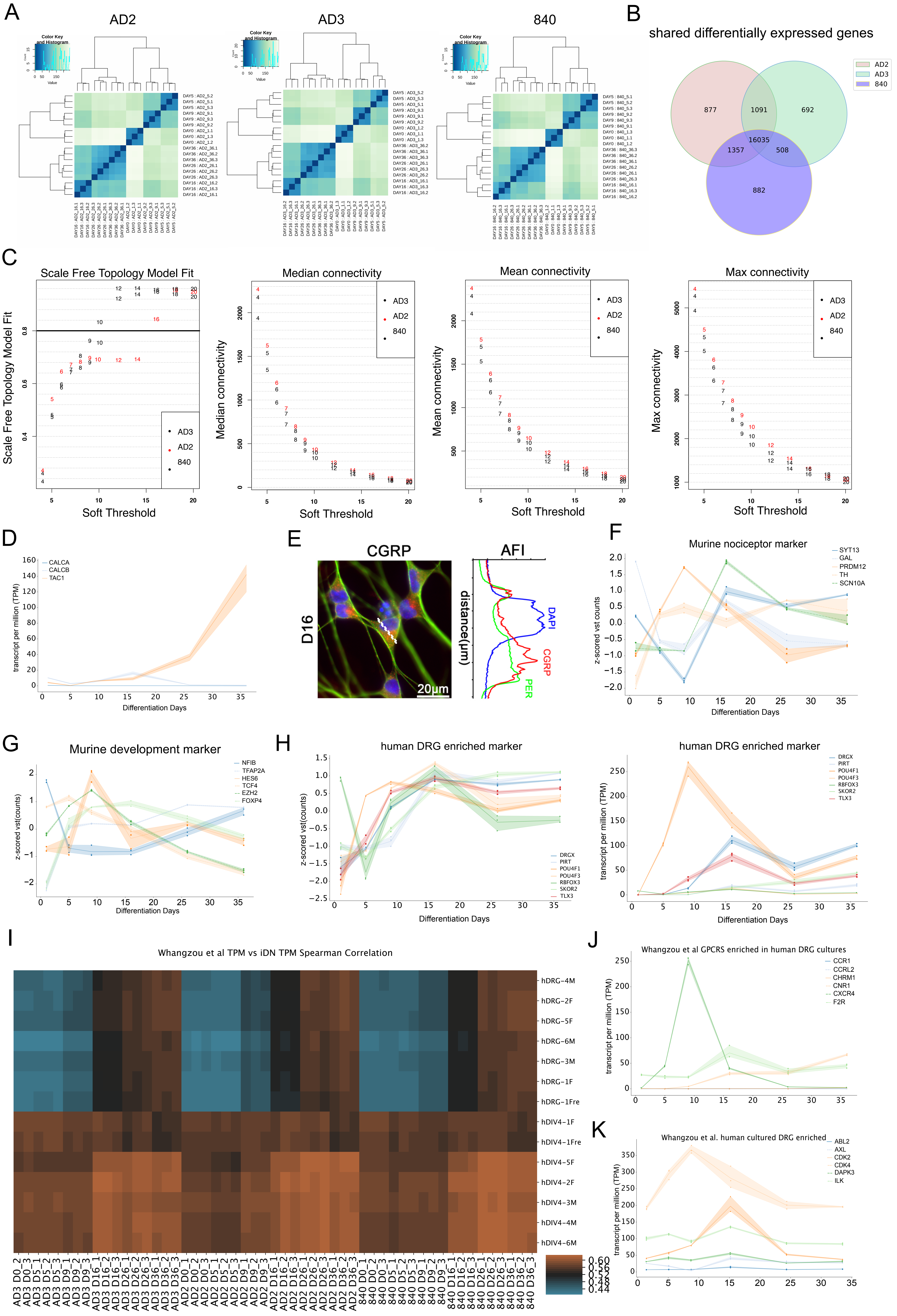

### Supplementary Figure 2

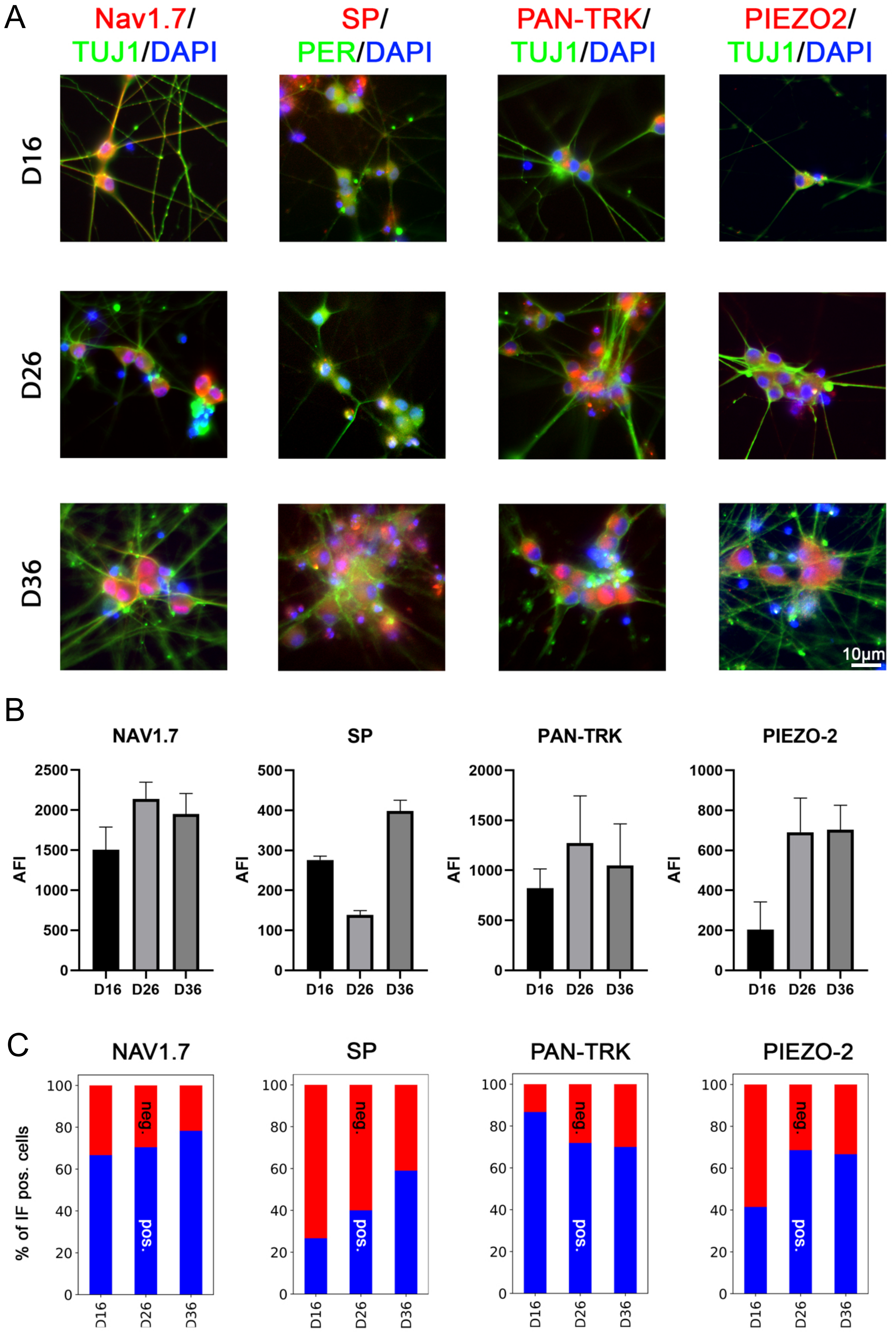

### Supplementary Figure 3

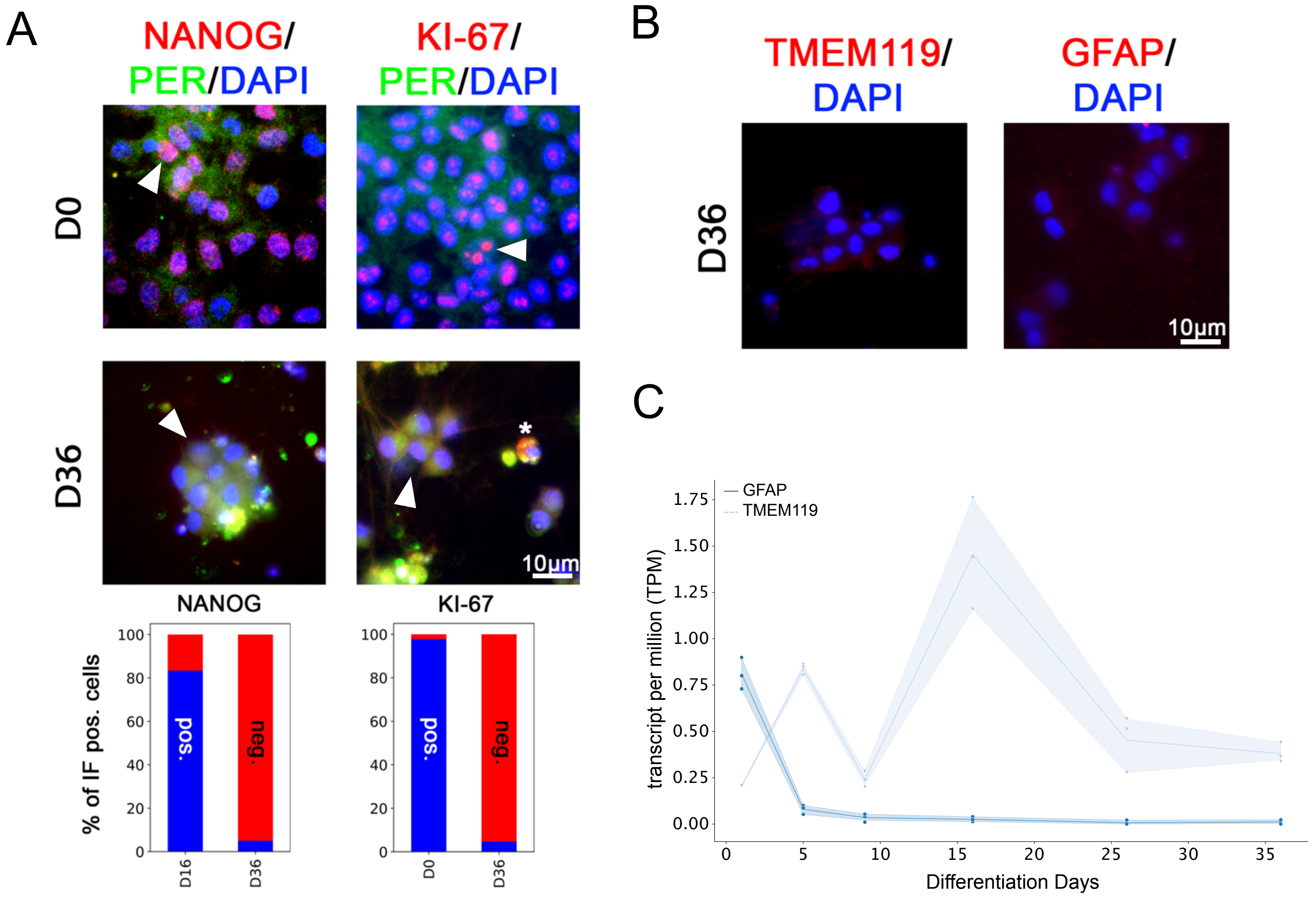

### Supplementary Figure 4

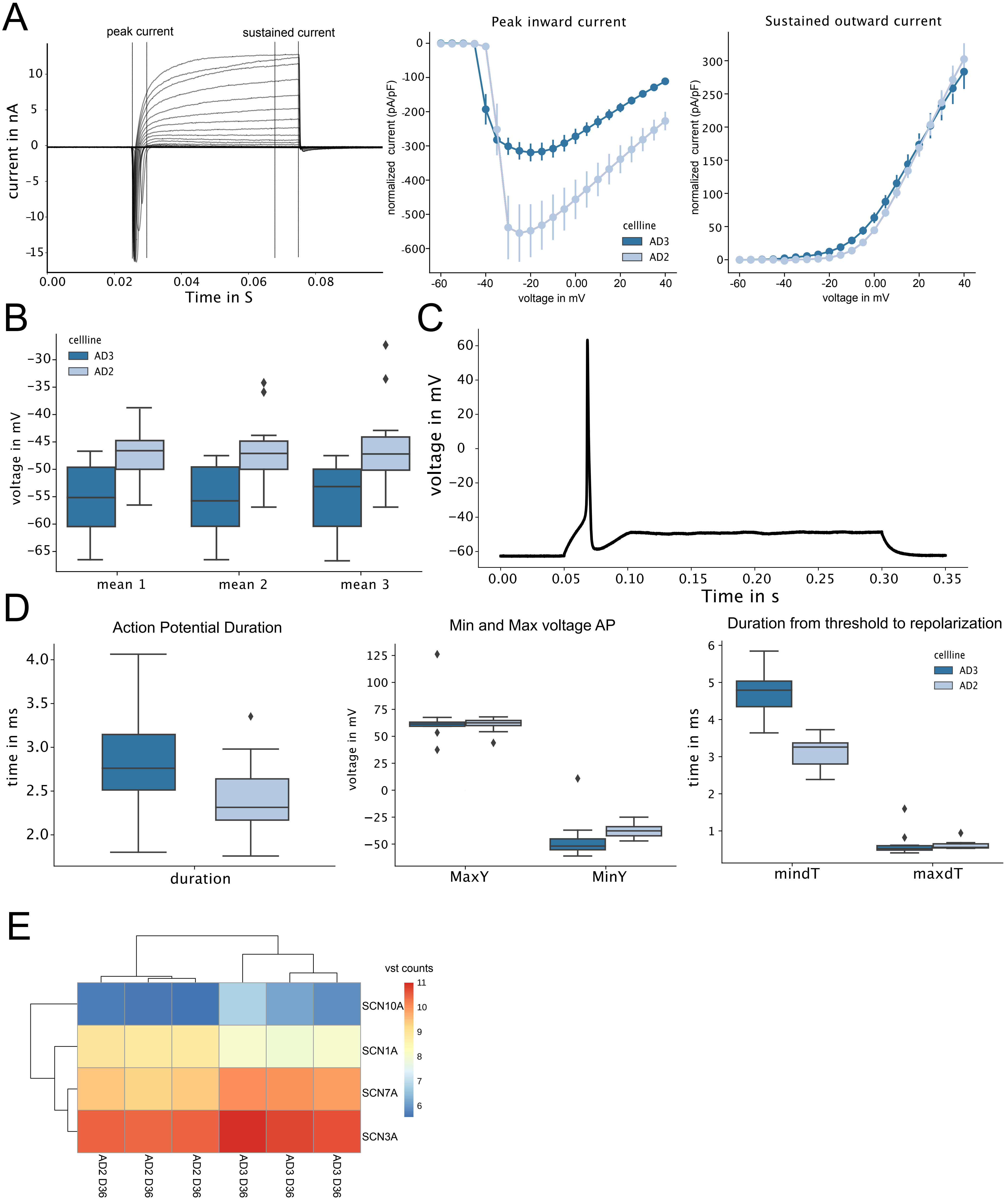

### Supplementary Figure 5

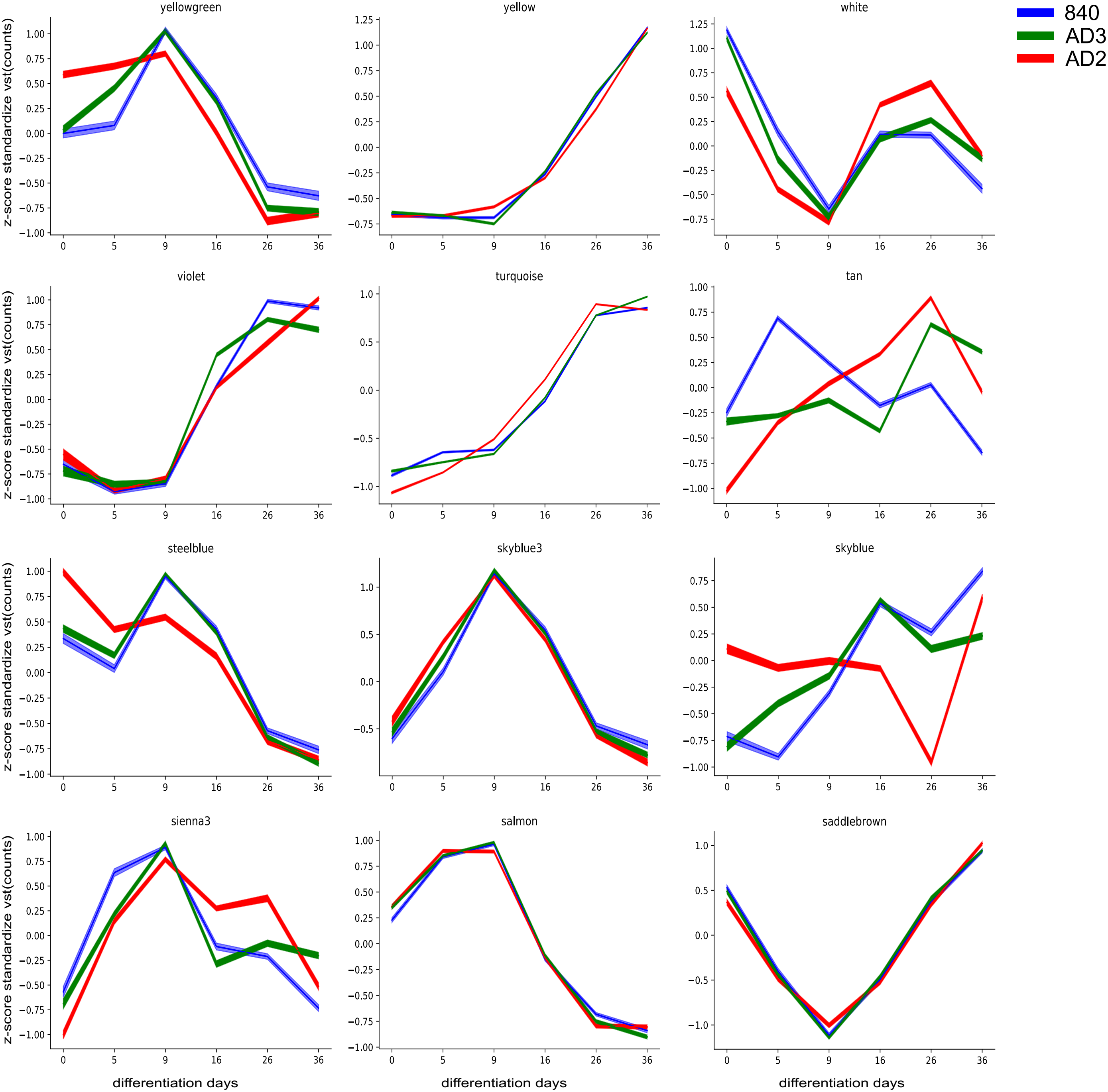

### Supplementary Figure 6

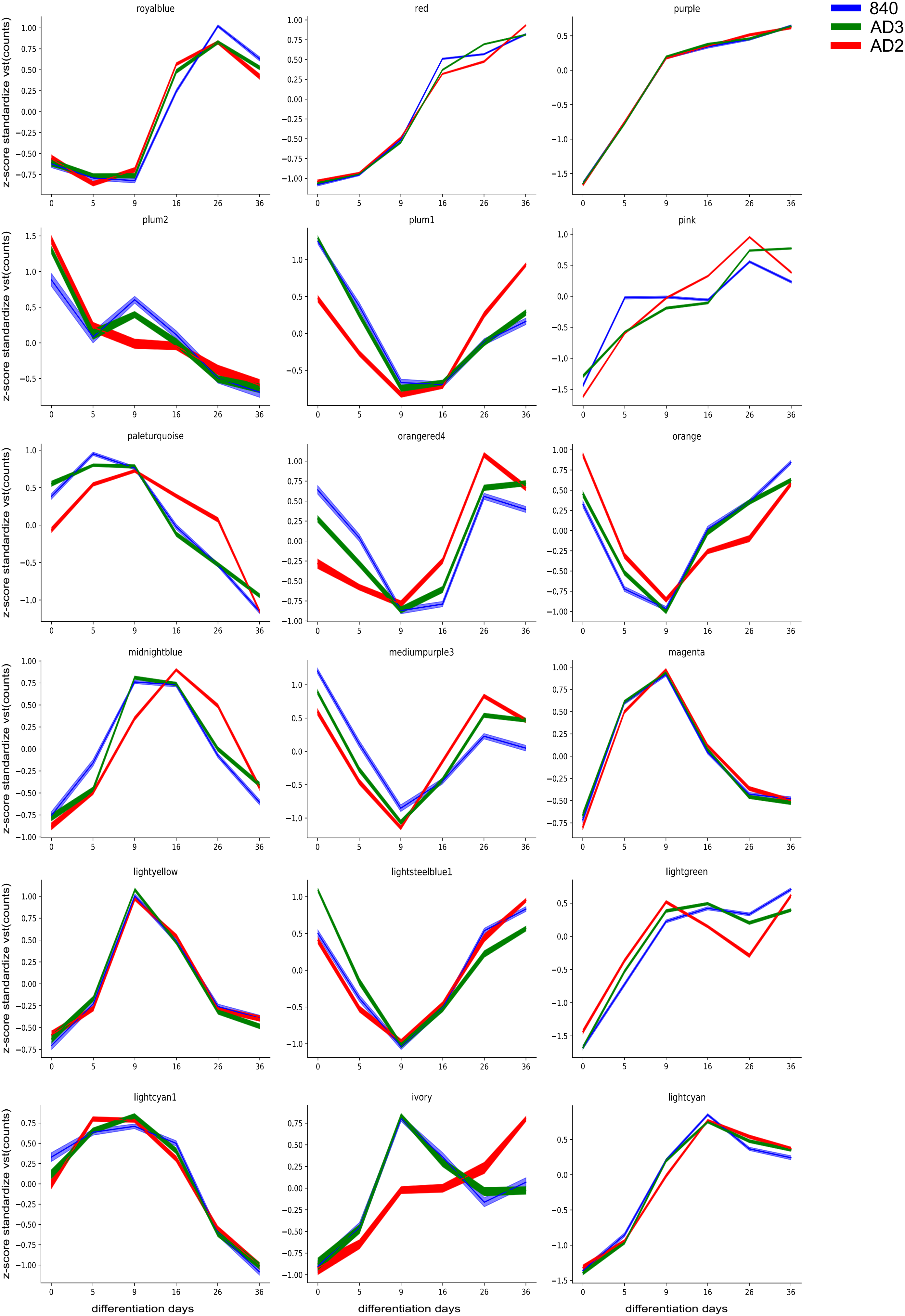

### Supplementary Figure 7

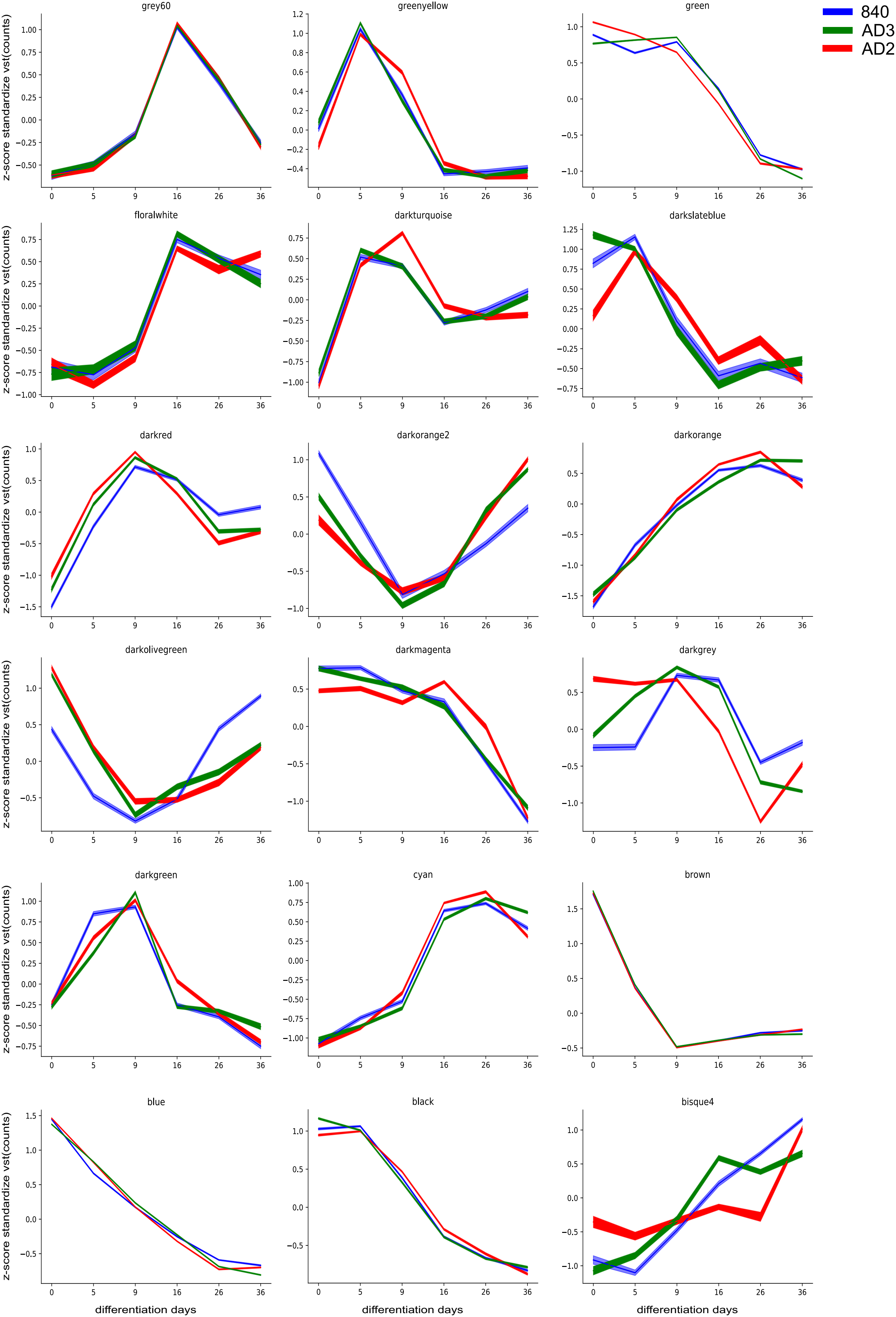

### Supplementary Figure 8

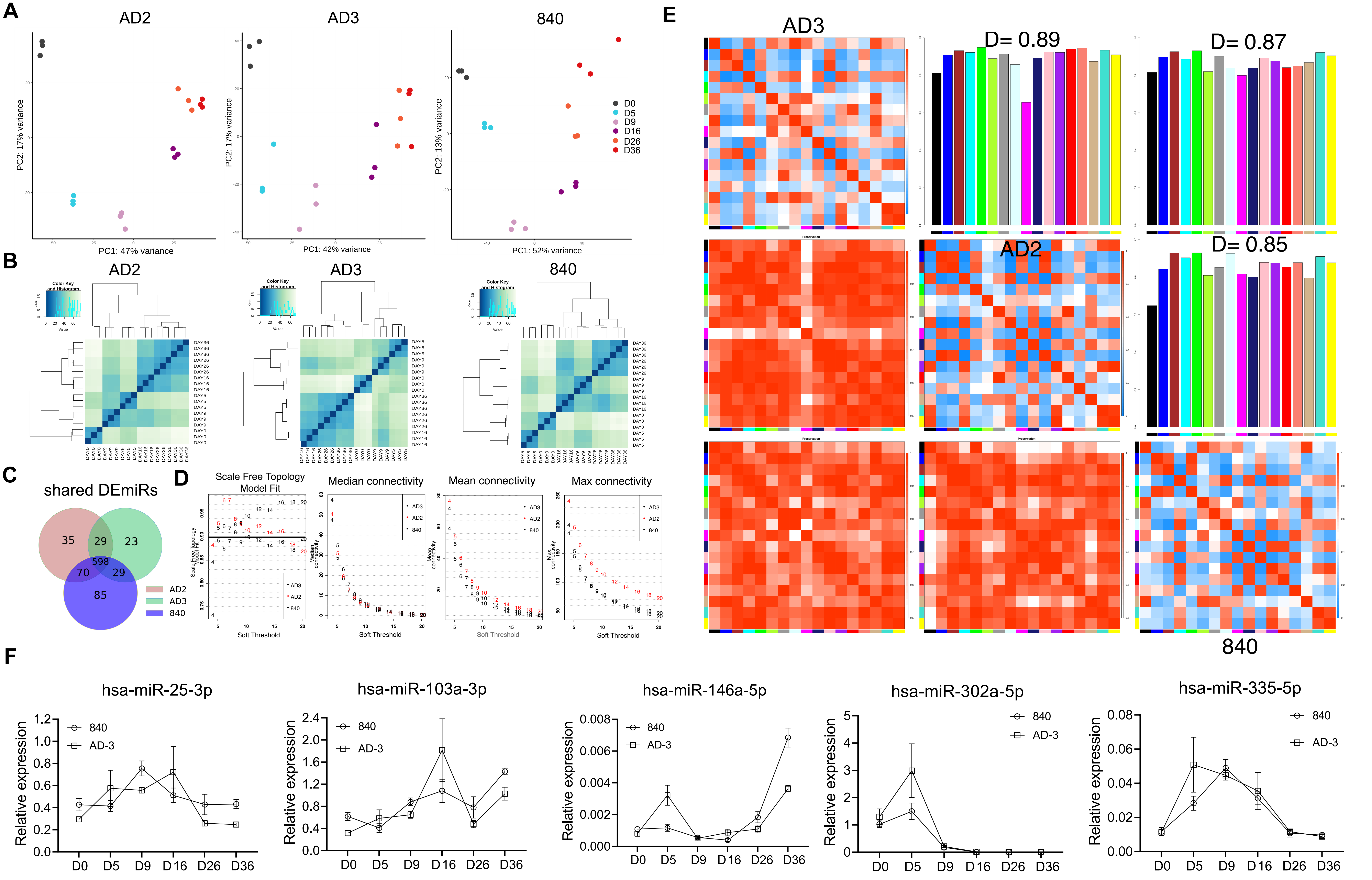

### Supplementary Figure 9

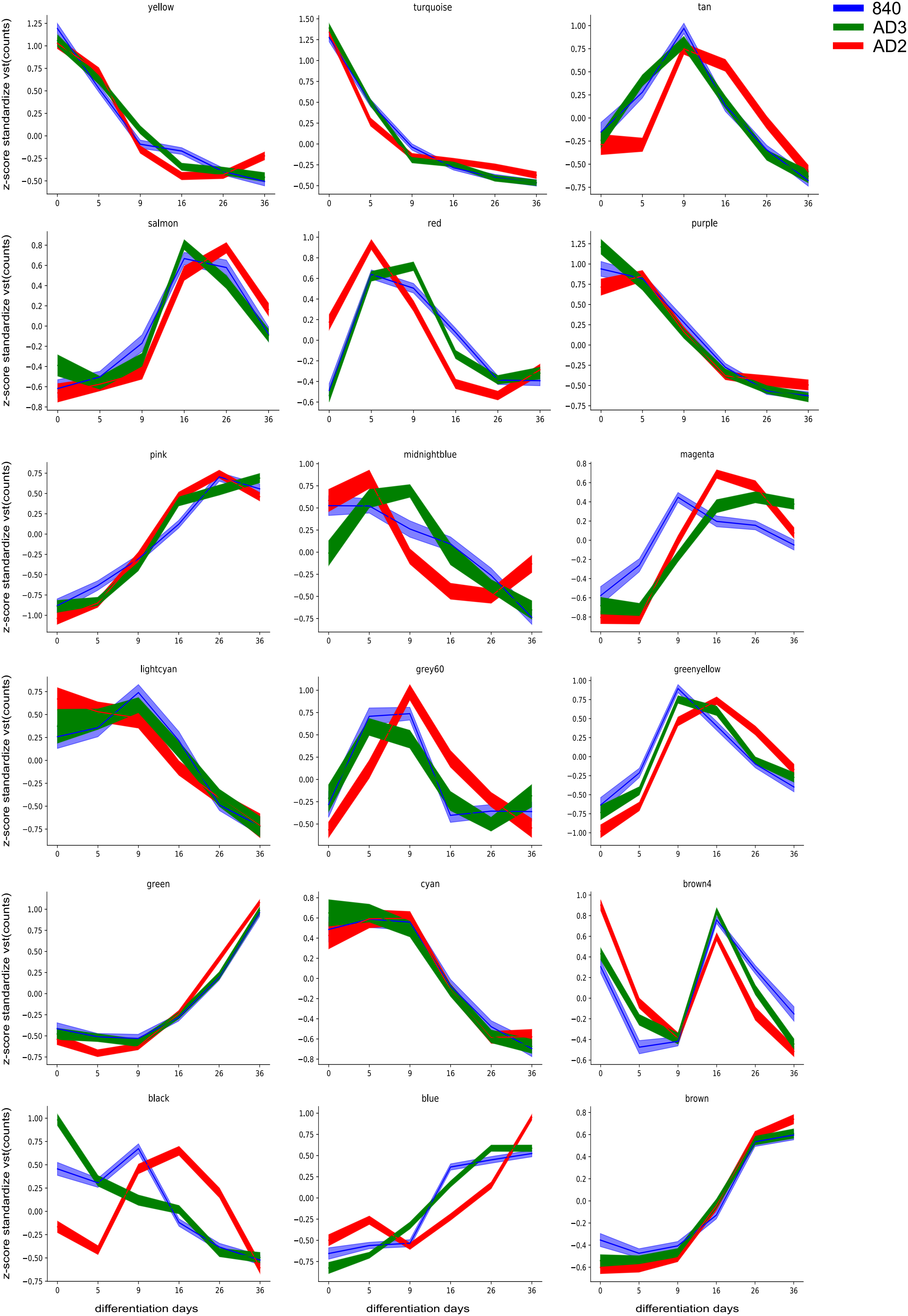

### Supplementary Figure 10

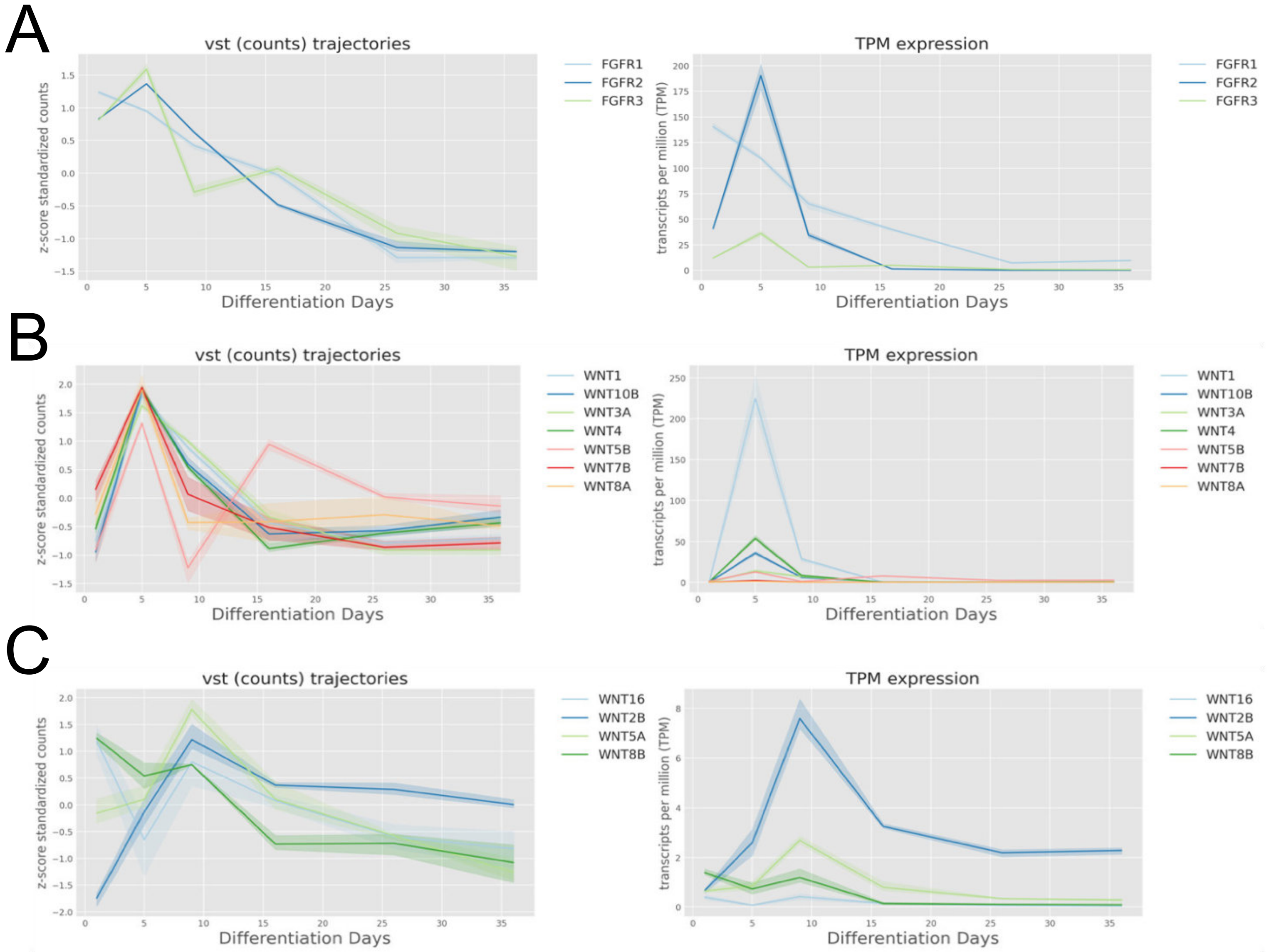
